# Reliable Evaluation of Transductive Population Graph Neural Networks for Multisite Autism fMRI

**DOI:** 10.64898/2026.09.15.751719

**Authors:** Keliang Wan, Zongqing Chen, Guixian Liu, Biantian Yu, Qian Zhang, Fanghong Zhang, Ning Zhong, Hongzhi Kuai

**Affiliations:** School of Mathematical Sciences, Chongqing Normal University, Chongqing, 401331, China; College of Artificial Intelligence and Advanced Interdisciplinary Sciences, Chongqing Normal University, Chongqing, 401331, China; Chongqing Key Laboratory of Brain-Inspired Cognitive Computing and Educational Rehabilitation for Children with Special Needs, Chongqing Normal University, Chongqing, 401331, China; School of Science, Tianjin Chengjian University, Tianjin, 300384, China; Department of Child Health Care, Children’s Hospital of Chongqing Medical University, Chongqing, 400014, China; National Clinical Research Center for Child Health and Disorders, Ministry of Education Key Laboratory of Child Development and Disorders, Chongqing Key Laboratory of Child Health and Nutrition, Chongqing, 400014, China; National Center for Applied Mathematics, Chongqing Normal University, Chongqing, 401331, China; Faculty of Engineering, Maebashi Institute of Technology, Gunma, 371-0816, Japan; School of Artificial Intelligence, and the CMEC Key Laboratory of Intelligence Health, Chongqing University of Posts and Telecommunications, Chongqing, 400065, China; Chongqing Key Laboratory of Computational Intelligence, Key Laboratory of Cyberspace Big Data Intelligent, Security, Ministry of Education, and Key Laboratory of Big Data Intelligent Computing, Chongqing University of Posts and Telecommunications, Chongqing, 400065, China

**Keywords:** graph neural networks, autism, multisite fMRI, transductive learning, unseen-site evaluation, reproducibility

## Abstract

Reliable population-graph evaluation requires separating cohort-conditioned discrimination from performance at an unseen site. We evaluate a recovered site-aware graph neural network in ABIDE-I (871 participants, 20 sites), distinguishing archived intervention results from newly recomputed controls. Archived cohort AUC was approximately 0.94, with high discrimination retained by site-only connectivity. A reconstructed ten-fold empirical site-prior classifier achieved pooled out-of-fold AUC 0.506 (95% site-resampling interval [0.418, 0.546]); simple site proportions therefore did not reproduce the high cohort result. In a budget-matched support-location experiment, the across-site mean AUC difference was +0.252 [0.184, 0.322] with directed edges but −0.003 [−0.020, 0.015] after symmetrisation, demonstrating implementation sensitivity rather than a stable support-location advantage. A supplementary leave-one-site-out analysis compared scoring-time population-edge access after the same source-site training procedure. Mean site-macro AUC was 0.577 with access and 0.580 without it; the paired difference was −0.0037 [−0.0179, 0.0111]. These observations do not establish equivalence or a complete mechanism. Explicit access contracts, compatible estimators and run-level provenance are necessary to interpret high transductive cohort performance.

## 1. Introduction

Medical-imaging models are often developed under acquisition conditions and patient distributions that differ from those encountered in use. Domain shift therefore makes out-of-domain evaluation central to reliable prediction. Recent medical-imaging work has examined reliability and generalization across heterogeneous domains [1, 2]. In multisite resting-state functional MRI (rs-fMRI), acquisition site is associated with scanner hardware, protocols, recruitment, and preprocessing variation [3, 4, 5].

Population graph neural networks introduce a further distinction. They connect subjects through imaging similarity and non-imaging phenotypes, allowing predictions to use cohort context [6, 7]. In transductive evaluation, training and test subjects share a graph: test labels are excluded from fitting and selection, but test imaging and phenotypes can influence graph construction and message passing. This information-access regime is distinct from direct label leakage [8]. Its performance estimate may nevertheless depend on information unavailable for an unseen acquisition site or a prospective single-subject prediction.

Recent population-graph studies integrate site-specific and site-invariant information in multisite autism and psychiatric imaging [9, 10]. Alongside improving prediction, this raises an evaluation question: which information pathways sustain high cohort discrimination, and do they support discrimination at an unseen site? A recovered implementation motivating this study achieved ABIDE-I cohort AUCs of approximately 0.90–0.96, substantially above values commonly reported under controlled multisite evaluation [11, 12]. Understanding that discrepancy requires interventions on the evaluated system rather than another performance comparison alone.

We examine one site-aware heterogeneous population-GNN using a numerically matched re-implementation and a fixed cohort. The contribution is an evaluation study, rather than a new diagnostic architecture. We define the information-access contract, intervene on graph sources, features and supervision, compare architecture-matched heads, and examine cross-pipeline and site-held-out results. A paired support-location experiment additionally tests sensitivity to edge construction and support composition. The revised analysis also includes a training-only site-prior baseline and a supplementary scoring-time LOSO access comparison. Archived and newly recomputed runs are kept separate; numerical proximity is not treated as proof of run identity. The resulting evaluation strategy separates observed performance dependencies from mechanistic attribution and distinguishes cohort-conditioned discrimination from unseen-site generalization.

## 2. Related work

### 2.1. Graph-based neuroimaging and multisite learning

Population graphs pool information across subjects using imaging and phenotype relations [6, 7, 13]. Applications include multisite autism classification [14, 15, 16] and federated models that combine site-specific and shared representations. SFPGCL addresses multisite ASD identification [9], while FGDN uses federated population graphs for major-depression diagnosis [10]. Standard graph convolution, attention and neighbourhood aggregation provide the relevant message-passing operators [17, 18, 19].

Other graph and multimodal approaches use different units of analysis. Saponaro et al. combined structural and functional MRI for ASD classification with nested cross-validation on ABIDE-I/II [20]. CAS-GNN combines individualized multimodal brain graphs with adversarial and supervised contrastive site alignment and evaluates four major ABIDE-I sites under leave-one-site-out testing [21]. Those graphs connect regions within a subject; the population graph examined here connects subjects. Their cross-site results therefore address related questions under different information-access regimes.

### 2.2. Domain shift and evaluation reliability

Connectome-based performance varies with representation, preprocessing, model and validation protocol [22, 11, 23]. Site and scanner structure motivates harmonization and domain-aware testing [24, 5, 25]. Reliability and adaptation studies in medical imaging likewise distinguish in-domain accuracy from performance across acquisition domains [1, 2]. In Alzheimer’s disease imaging, Akhavan Aghdam et al. combined a survey with open-source model evaluation, separating reproduction on the original data from generalization across ADNI and OASIS [26]. For population graphs, evaluation must additionally specify which target nodes, phenotypes and edges are visible. Medical-imaging models can exploit dataset-specific regularities rather than the intended biological signal [27, 28, 29]. Leakage, optimistic validation and small evaluation samples further complicate interpretation [30, 31]. Recommendations for neuroimaging prediction emphasize explicit validation and separation of exploratory from confirmatory analyses [32]. Our study complements this work by intervening on site-defined connectivity, train–test pathways, supervision and population heads within a specified transductive setting. It assesses the information underlying a performance estimate while treating unseen-site evaluation as a separate target.

## 3. Methods

### 3.1. Cohort and information-access regimes

The canonical ABIDE-I cohort [33] comprises 871 subjects (403 ASD and 468 typically developing) from 20 sites. We used C-PAC and NIAK preprocessing (filt_noglobal), AAL-116 and CC200 atlases, and HO-111 on C-PAC. Functional-connectivity correlations were Fisher-*z* transformed. Unless otherwise stated, reported reference-model results use AAL-116. Fitted preprocessing is fold-local: StandardScaler and PCA are fitted on optimization-training subjects, with held-out data transformed using the fitted quantities.

In the cohort protocol, labels define pre-generated stratified folds; test labels subsequently contribute only to scoring, not to optimization, preprocessing or model selection. Test imaging, site and sex remain visible to graph construction, and train–test and test–test edges are retained (Table 1). This defines the transductive information-access regime.

**Table 1:** Information-access contract (per outer fold).

| Information | Train | Validation | Test |
| --- | --- | --- | --- |
| Imaging features | ✓ | ✓ | ✓ |
| Site / sex | ✓ | ✓ | ✓ |
| Labels | ✓ | model selection only | **sealed** |
| Train–test edges | ✓ | ✓ | ✓ (transductive) |
| Test–test edges | ✓ | ✓ | ✓ (transductive) |
| Scaler / PCA fitting | fit on train only | transform | transform |

For leave-one-site-out (LOSO) evaluation, the target site is excluded from optimization, validation, early stopping, and fitted preprocessing. Scaler/PCA and the affinity-normalization reference use non-target training subjects. For the supplementary analysis reported here, source subjects alone enter optimization and validation. At scoring, target subjects are attached through cross-site same-sex edges in the access condition; the comparison condition omits those edges. Target-to-target population edges are absent in both conditions. LOSO therefore assesses unseen-site discrimination under this attachment rule. Relative to cohort evaluation, it jointly changes supervision availability, target-site topology and training-domain composition; it neither isolates those changes nor estimates every prospective deployment setting.

### 3.2. Evaluated reference model

We refer to the evaluated site-aware population graph neural network as SA-PopGNN. This descriptive name denotes the recovered reference architecture and its numerically matched re-implementation, rather than a newly proposed model. It combines a subject-level connectome encoder with a population head whose two channels distinguish same-sex from different-sex relations. Site metadata enter through edge construction, not as a diagnostic label.

The subject encoder has three GCN layers (141 → 64 → 64 → 32), SAGPool and DiffPool-style fusion with ChebConv scoring (*K* = 3). Each AAL-116 connectome is combined with atlas-derived context features into a 116 × 141 tensor. Subject-level connectome edges retain |*r*| > 0.5, carry weight |*r*|, omit self-loops, and aggregate in both directions after pairs are enumerated once with *i* < *j*. The encoder produces a 32-dimensional subject embedding.

Population affinity is A_i j_ = Σ_a∈{site,sex}_ 1[a_i_ = a_j_]. It is column-standardized using training rows as reference; values above 1.0 select edges. Sex agreement partitions selected edges into *E*_same_ and *E*_diff_. Affinity magnitude is not a numerical message weight: the population `TransformerConv` layers have no edge-feature dimension. The two relation channels use 32 → 32 transforms with one attention head per channel, combined as

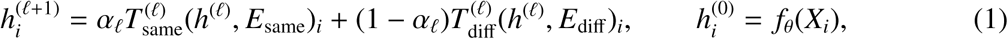

with learned channel weight α_ℓ_ and class logits MLP(*h^(L)^_i_*).

The independently written harness was checked against the recovered model under strict checkpoint loading. Numerical agreement establishes fidelity of that tested path, not recovery of every historical training artifact. The original saved inner-validation masks are unavailable. Reconstructed masks reproduce the supplied outer fold identifiers, but this does not independently establish the identity of every historical fit. The recovered implementation is treated as a case study; no claim of a new architecture or a complete reconstruction of its historical development is made.

### 3.3. Training and controlled comparisons

Outer folds use shuffled 10-fold `StratifiedKFold` with the fixed split seed. Each outer-training partition reserves 10% through `StratifiedShuffleSplit` for validation, using the same seed. Training uses Adam (10^−3^), weight decay 5 × 10^−5^, label-smoothed cross-entropy (0.1), dropout 0.3, at most 400 epochs and patience 200. Selection maximizes validation accuracy plus AUC; test folds are never used for selection. StandardScaler and PCA(32) are fitted only on the optimization-training partition with fixed preprocessing seeds. GPU arithmetic may retain small nondeterminism, so reported values refer to the frozen artifacts rather than bit-level retraining guarantees.

#### Graph and feature controls

Holding the architecture and cohort protocol fixed, we evaluate site-only, sex-only, edge-count-matched random and subject-only/no-graph conditions. The random control redraws the sex+site edge count over *i* < *j* pairs without matching per-node degrees. Global label-permutation controls permute labels once per seed (random seed 1000 · seed + 7), retaining the original folds, features and population edges.

Four additional interventions use the site-only graph and seed 666: (A) constant node features; (B) within-site feature shuffling; (C) train–test edge removal; and (D) degree-preserving rewiring. They probe feature content, fused-feature assignment, cross-partition pathways and the specific connectivity pattern, respectively. The shuffle code reassigns fused features across all subjects within a site, potentially crossing training, validation and test roles, while retaining each subject’s original connectivity topology, site, sex and label. It is not a permutation of complete individual imaging inputs; the historical run’s exact permutation map was not retained. Rewiring uses `networkx.double_edge_swap`, with nswap = max(100, 5*m*) and max_tries = max(1000, 50*m*) for *m* edges, per fold. Random-forest and linear-SVM classifiers on upper-triangular AAL connectivity entries provide classical imaging-only references.

#### Architecture and supervision controls

Head ablations retain the subject encoder, sex+site edges, folds and training protocol while replacing the population head with single-channel Transformer-Conv, GAT, GCN or a no-graph MLP. A dual-channel control supplies identical edge sets to both branches. A separate topology-only Parisot-GCN-C comparator uses Chebyshev order three, imaging-similarity and sex/site-defined edges, and no site/sex edge features; a further variant adds site as a node feature. Matched data and folds do not imply matched optimization effort across architectures.

The site-wise supervision-masking protocol excludes each target site’s subjects from both the loss and validation/model selection while retaining their graph nodes and message passing. Each target subject is evaluated once. Its comparison with cohort 10-fold results is descriptive because the training partition also changes. A separate half-site experiment has incomplete split provenance and is confined to Supplementary Note S2.

### 3.4. Paired support-location and sensitivity experiments

The paired experiment fixes graph visibility and supervision budget within each target-site/fold cell (Figure 6). Let *T* be target-site test subjects, *S* target-site support, *O* other-site trainable subjects and *V* fixed validation. Draw *R* ⊂ *O* to match *S* in total size and class counts, and set *C* = *O* \ *R*. Training labels come from *C* ∪ *S* in ON and *C* ∪ *R* in OFF; the other support subjects remain graph-visible. Graph, validation, initialization and training schedule are identical within each pair. Preprocessing is fitted on *S* ∪ *O*; *T* and *V* are transform-only. *V* has no population edges and its labels serve model selection only.

Same-site population edges connect eligible nodes in *T* ∪ *S* ∪ *R* ∪ *C*. The primary construction records each pair once (*i* < *j*), so message direction follows subject index. The second TransformerConv channel receives no edges but retains its root-weight transform; this path adds no graph self-loops or row normalization. It is distinct from the Parisot comparator’s row-normalized Chebyshev propagation and from the separately built main-model/head-ablation graphs. We repeat the paired experiment with both edge directions included, leaving all other elements unchanged. A six-pair orientation diagnostic is documented in Supplementary Note S5.

The analysis comprises 18 eligible sites, 88 site–fold cells and three fixed training seeds (666, 2025 and 2026). SDSU contributes three folds; other included sites contribute five. CALTECH and CMU have no eligible cells. A separate, pre-declared sensitivity analysis repeats support draws at LEUVEN_1, PITT and YALE using three fixed sampling seeds. Because changing *R* changes *C*, both arms’ supervised subject sets can change even though their graph and supervision budgets remain fixed. Neither budget matching nor support resampling separates location from subject identity and covariate composition.

#### Empirical site-prior baseline

For each outer test subject, the score is the ASD fraction among optimization-training subjects from that site. Validation labels are excluded; a fold-wide optimization-training prior is used if the site has no training subject. We reconstructed the seed-666 outer ten-fold split and inner validation split from the supplied recipe; outer identifiers match the archived OOF file. Every subject is scored once. The interval uses 10,000 site-resampling draws (seed 20261015), retaining repeated sites. This analysis uses labels for the training prior only, not test-score construction. A separate paired-protocol baseline uses *C* ∪ *S* for ON and *C* ∪ *R* for OFF and is not interchangeable with this cohort baseline (Supplementary Note S7).

#### Supplementary scoring-time LOSO access analysis

For each held-out site and training seed (666, 2025 and 2026), the same source-site optimization and validation sets are used in both conditions; the split seed is 666. StandardScaler and PCA(32) are fitted on source optimization-training inputs only. The supplied driver trains on the source graph with Adam, learning rate 10^−3^, weight decay 5 × 10^−5^, label smoothing 0.1, at most 300 epochs and patience 150. Selection uses source validation accuracy plus AUC. The access condition adds cross-site same-sex population edges only at scoring; the no-access condition omits this insertion. The training graph and objective do not differ by condition. The driver executes training separately with matched seeds, but these are not two training-time graph interventions. Target imaging is encoded separately from source imaging and the embeddings are concatenated for population-head scoring. Both conditions omit target–target population edges. Assertions reported by the execution campaign cover source supervision and target-edge counts; the supplied predictions independently support the endpoint recomputation, not a complete reconstruction of all external model dependencies.

#### Evidence scope

The later complete-imaging-swap campaign is excluded from the manuscript’s mechanism evidence because its graph protocol did not correspond to the original cohort experiment and its observed perturbation response was not localized. A separate added-access endpoint-swap campaign is also excluded from the main evidence chain; two seeds deviated from the planned split. These exclusions do not invalidate or validate the earlier archived controls. No inference about leakage or an abnormal encoder pathway is made from unlocalized logit changes.

### 3.5. Statistical analysis and reproducibility

#### Cohort and site-held-out endpoints

The primary cohort endpoint is pooled out-of-fold AUC: one ROC-AUC over all held-out subjects, using softmax(*z*)[:, 1] (equivalently the ranking of *z*_1_ − *z*_0_). Mean-fold AUC assigns equal fold weights; site-macro AUC assigns equal site weights. These estimators are reported separately. For the supplementary LOSO access analysis, the primary endpoint is the equal-weight mean site AUC, computed separately for each training seed and then averaged. Pooled AUC is secondary; predictions from different seeds are never stacked as independent subjects. Older LOSO summaries are retained only as historical records in Supplementary Table S5. Mean-fold reference values are recomputed from frozen folds joined by subject ID; historical values not reproduced from released predictions are excluded.

For cohort estimates, site-clustered bootstrap resamples the 20 sites with replacement (1,500 replicates); subject-level bootstrap intervals are also available. These procedures resample saved predictions without refitting or rebuilding the graph. Their uncertainty is conditional on fitted predictions and omits full retraining variability; shared folds and cross-site edges may induce dependence beyond site clustering.

The historical 13-contrast analysis is retained for traceability in Supplementary Notes S1 and S6; its original *p*-values and Holm adjustments are not endorsed as current inferential evidence. Table 5 instead reports effect sizes and paired site-bootstrap intervals recomputed from a separate set of control-run OOF archives (10,000 draws, seed 20261015). Subject identifiers, labels and fold identifiers match within each comparison. The control Full AUC is 0.949842, whereas a separately supplied main-model OOF archive yields 0.939609. Neither is silently substituted for the manuscript’s archived 0.9405: configuration and historical run identity remain unresolved. These new intervals do not validate the historical *p*-values or recover the original comparison family.

For the supplementary LOSO contrast, the same resampled sites are used for both conditions and all three seeds (5,000 draws, seed 20261016). Site differences are averaged within each seed and then equally across the three seeds. Intervals are conditional on these fitted predictions and fixed seeds. Bootstrap positive-draw fractions are distribution descriptions, not *p*-values. Marginal 95% intervals are not simultaneous intervals or equivalence tests.

#### Paired support-location endpoint

For each training seed we average ON−OFF AUC over eligible folds within each site, then average equally over sites and the three fixed seeds. This across-site mean ΔAUC is the archived population-macro endpoint; a within-site fold mean is called a site effect here. This endpoint was implemented after training and is treated as a post-hoc analysis; it was not pre-registered. Its interval resamples sites (10,000 replicates), with an exact two-sided sign test over site effects and a null-centred bootstrap *p* using the finite-simulation floor 1/(*B* + 1). Per-site intervals condition on fitted models and fixed seeds. Per-site *p*-values and Holm adjustments based on the across-site effect distribution are withdrawn; that distribution is not a valid sampling unit for an individual site. Support-draw sensitivity is descriptive and is not pooled into this endpoint.

The frozen registry distinguishes cohort, site-wise and architecture-control protocols. Demographic, calibration, sex-stratified, label-proximity and attribution analyses are secondary; their scope is stated in the supplement. Figures 6–7 are rendered from frozen inputs using a released Python/Matplotlib script; they contain no newly inferred values. ChatGPT/Codex (OpenAI) assisted editorial revision and figure-code preparation under author review; the specific model version for that code assistance was not retained. Plotting inputs, script and dependency version are supplied for reproducibility.

## 4. Results

### 4.1. Implementation fidelity and cohort replication

Figure 1 summarizes the pipeline; Supplementary Figures S4–S7 give the full study-design panels. Under strict checkpoint loading, the re-implementation matched recovered model outputs to maximum absolute difference 7 × 10^−7^. The Parisot harness also reproduced its independent reference transductive AUC of 0.665. The archived manuscript reports SA-PopGNN cohort AUC 0.939 ± 0.002 across three seeds (0.941, 0.937 and 0.940; mean ± SD). These are historical campaign summaries. The separately supplied seed-666 main-model OOF file recomputes to 0.939609 (mean-fold AUC 0.942460), and is not assumed to be the archived 0.9405 run merely because the values are close. Additional repetitions are reported in Supplementary Table S7; not every intervention was repeated across seeds.

**Figure 1:**
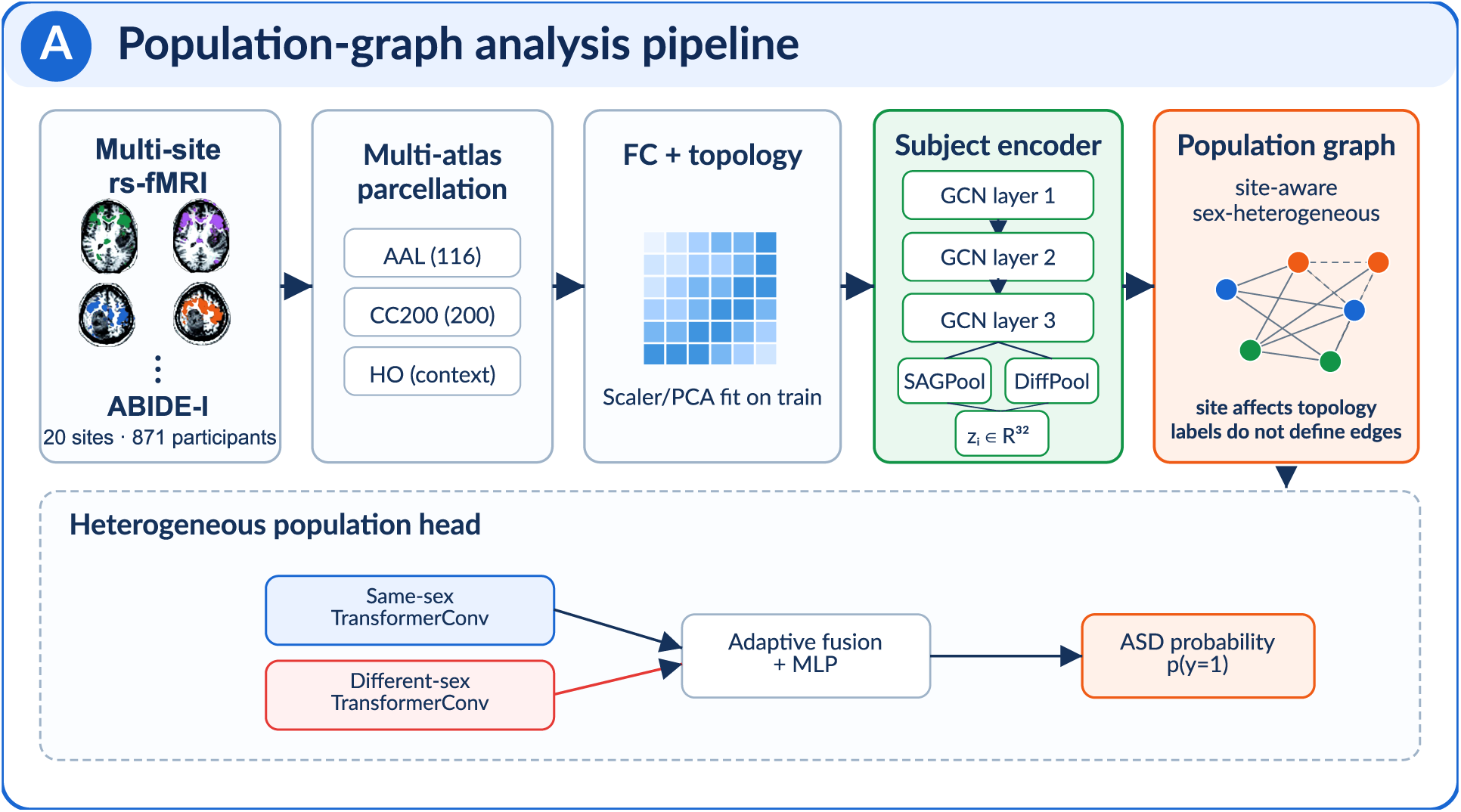
Analysis pipeline. Multisite ABIDE-I rs-fMRI undergoes multi-atlas parcellation, connectome construction and fold-local scaler/PCA fitting before subject encoding and population-graph prediction. Site/sex-defined edges feed separate same-sex and different-sex channels. The four brain icons are schematic site exemplars; the ellipsis represents additional sites in the 20-site cohort. Full-size evaluation panels and the composite overview appear in Supplementary Figures S4–S7. **AI-assisted illustration:** ChatGPT image-generation tools (OpenAI; accessed September–October 2026, model version not retained) assisted initial drafting. The authors reconstructed and verified the final vector artwork and scientific content.

### 4.2. Graph sources and demographic controls

Archived full-model pooled OOF AUC was 0.941 (site-clustered interval 0.867–0.976), while site-only edges retained 0.948 (0.875–0.982). Sex-only, random and subject-only conditions yielded 0.565, 0.587 and 0.587, respectively (Table 2; Figure 2). These archived point estimates and intervals are distinguished from the separately supplied control-run predictions in Table 5; the historical significance tests are not used to support current conclusions.

**Figure 2:**
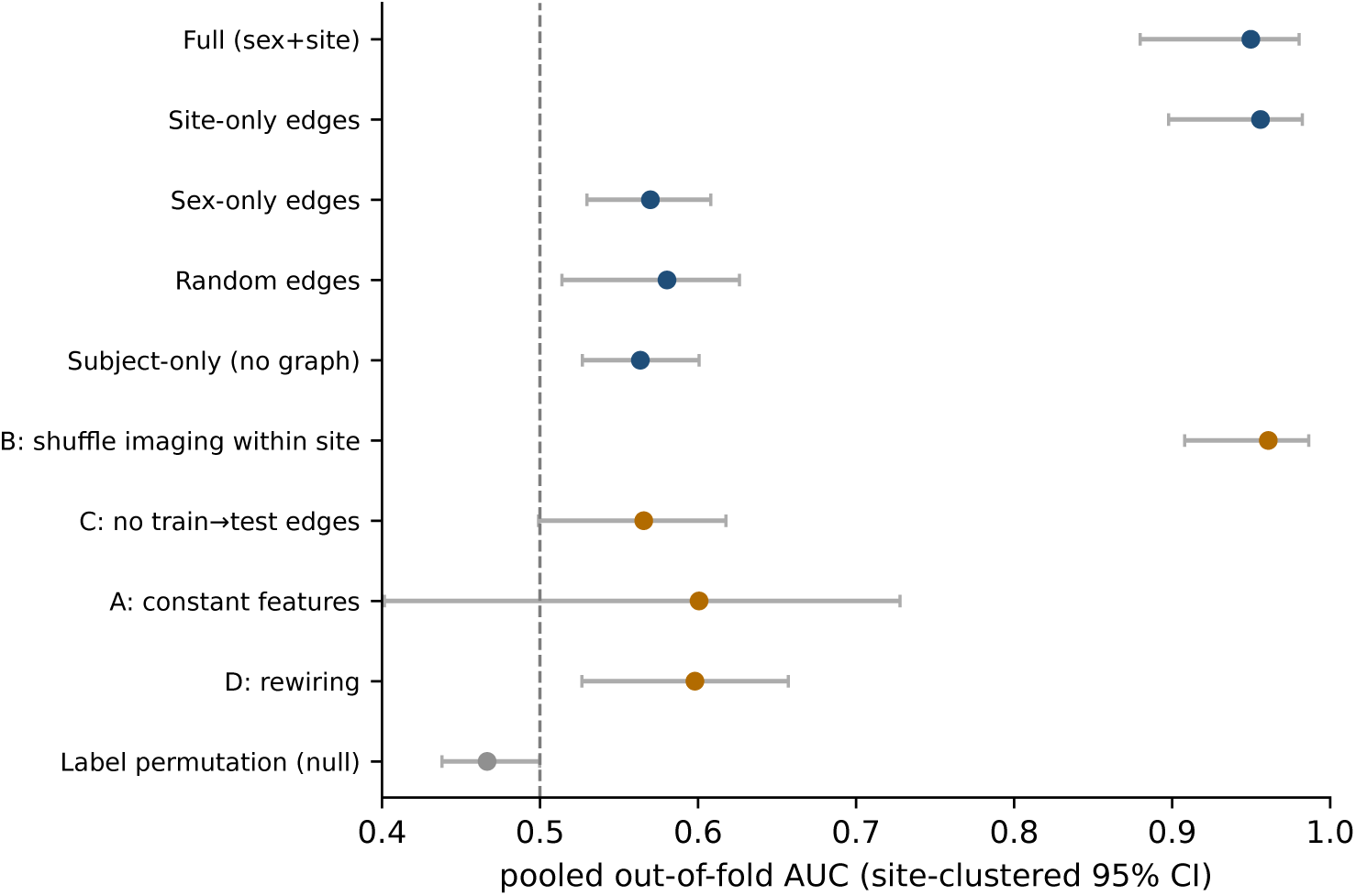
Archived cohort summaries (historical fitted-run linkage remains limited): pooled out-of-fold AUC by population-edge setting and controlled intervention, with site-clustered 95% CI (dashed line = 0.5 reference).

**Table 2:** Archived cohort intervention summaries, canonical ABIDE-I (871), seed 666 unless noted. Pooled OOF AUC and archived site-clustered intervals. Historical fitted-run identity is not resolved by numerical proximity; these rows are not the newly recomputed control runs in Table 5.

| Setting | Pooled OOF AUC (site-clustered 95% CI) |
| --- | --- |
| Full model (sex+site) | 0.941 (0.867–0.976) |
| seeds 2025 / 2026 | 0.937 / 0.940 |
| Site-only edges | 0.948 (0.875–0.982) |
| Sex-only edges | 0.565 (0.518–0.600) |
| Random edges (count-matched) | 0.587 (0.532–0.633) |
| Subject-only (no graph) | 0.587 (0.542–0.627) |
| *Controlled interventions (site-only graph)* |  |
| B: shuffle features within site | 0.960 (0.903–0.985) |
| C: no train–test edges | 0.562 (0.498–0.606) |
| A: constant node features | 0.600 (0.438–0.699) |
| D: degree-preserving rewiring | 0.597 (0.528–0.653) |
| Illustrative label-permutation controls (s666 / s777) | 0.492 / 0.464 |

Two illustrative global label-permutation controls yielded AUCs of 0.492 and 0.464. They did not reveal gross label contamination in those runs, but cannot exclude all leakage or provenance errors. Site as a node feature gave 0.504, and the cohort site×diagnosis association was modest (Cramér’s *V* ≈ 0.16; χ^2^ *p* = 0.225). A regularized demographic logistic-regression baseline likewise failed to reproduce the cohort AUC: site+sex achieved 0.5131 (site-clustered interval 0.425–0.555), and adding their interaction gave 0.5006. Across three seeds, pooled demographic AUCs differed by at most 0.011 (Supplementary Table S6). These controls constrain simple demographic explanations, not all site-typical imaging distributions. The newly recomputed empirical site-prior baseline scored all 871 subjects from all 20 sites, with no fallback, and achieved pooled OOF AUC 0.5056 (site-resampling 95% interval [0.4176, 0.5464]). This simple optimization-training site-proportion rule did not reproduce high cohort discrimination; it does not exclude nonlinear or other site-related pathways. Site sample sizes ranged from 11 to 172 (median 30.5); diagnosis counts by site are in Supplementary Table S1.

The NIAK cohort results followed the same pattern: full-model AUC 0.946, site-only 0.958 and subject-only 0.570 (Table 3). Thus high cohort discrimination and its dependence on graph setting were observed under both preprocessing pipelines.

**Table 3:** Cross-pipeline consistency, *cohort 10-fold protocol only* (seed 666; transductive rows are pooled out-of-fold AUC and the imaging RF row is mean-fold AUC, as documented in Supplementary Material, Supplementary Table S4). Site-held-out (leave-one-site-out) values are reported separately in Supplementary Table S5 as historical records, not current evidence for a cohort-to-LOSO effect.

|  | C-PAC | NIAK |
| --- | --- | --- |
| Full model (sex+site) | 0.941 | 0.946 |
| Site-only | 0.948 | 0.958 |
| Subject-only (no graph) | 0.587 | 0.570 |
| No train→test edges | 0.562 | 0.593 |
| Imaging RF (10-fold) | 0.669 | 0.602 |

### 4.3. Feature and train–test pathway interventions

Archived within-site feature shuffling retained high cohort AUC (0.960, interval 0.903–0.985). This result concerns reassignment of fused features with original connectivity topology retained and possible movement across fold roles. It does not establish that the correspondence between a subject and their complete imaging inputs is unnecessary. Removing train–test edges reduced AUC to 0.562 (0.498–0.606); degree-preserving rewiring yielded 0.597 (0.528–0.653). Constant features yielded 0.600 (0.438–0.699), without clear evidence of above-chance discrimination (Table 2; Figure 3). These descriptive contrasts implicate graph connectivity and cross-partition pathways, without separately identifying an information-processing mechanism. NIAK train–test edge removal yielded 0.593.

**Figure 3:**
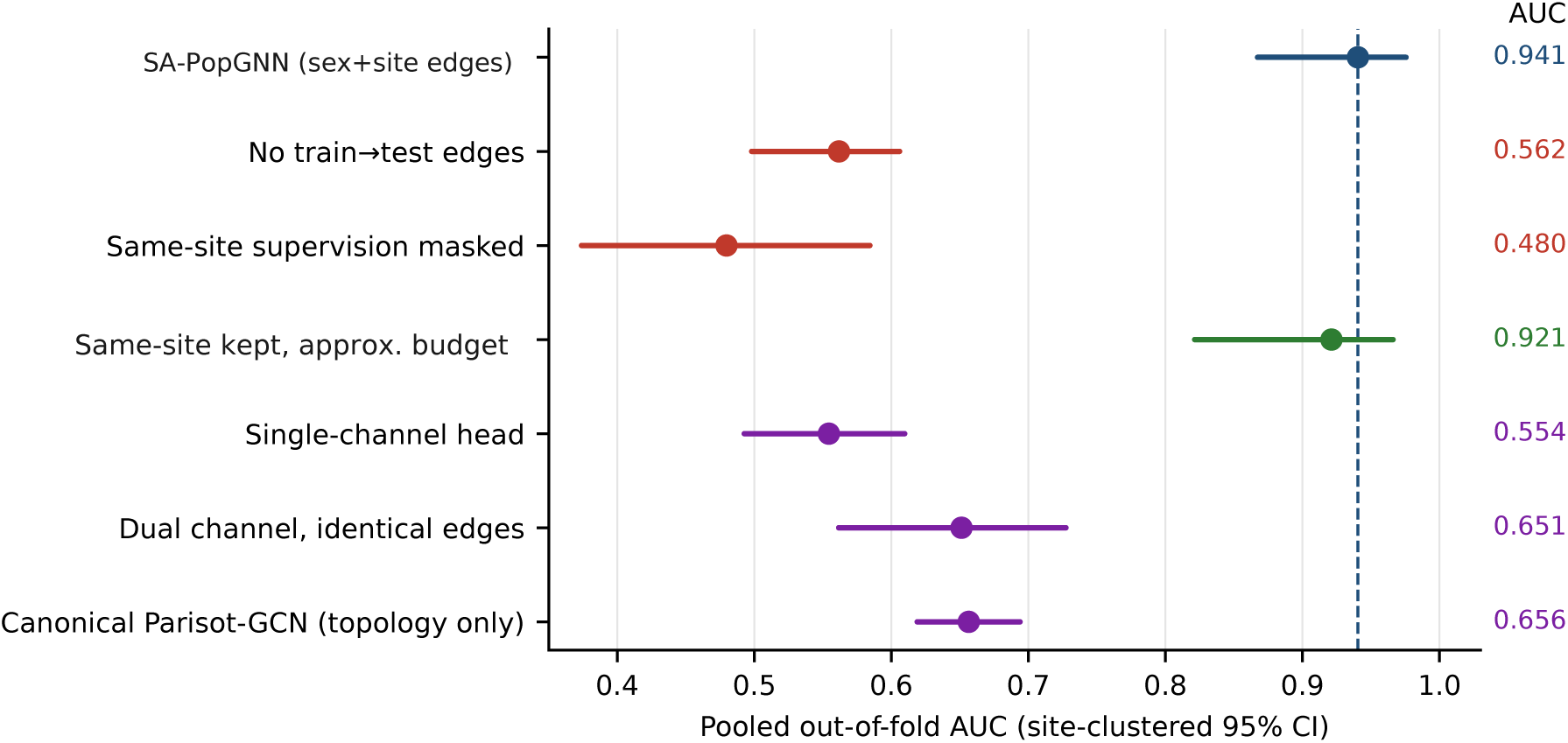
Archived controlled intervention summary; comparisons across protocols are descriptive. Pooled out-of-fold AUC (site-clustered 95% CI) for the full model and for each controlled intervention in intervention-ladder order: removing train→test edges, masking same-site supervision, the approximate supervision-budget control that retained same-site supervision (drawn under its historical experiment identifier, *budget matched*), and architecture-matched head replacements, with the canonical topology-only Parisot-GCN for reference. Every value is registry-backed; the dashed line is the full model. Side-by-side display does not imply a paired comparison across different protocols or evaluation subsets; the half-site budget arms are reported descriptively in the text.

**Figure 4:**
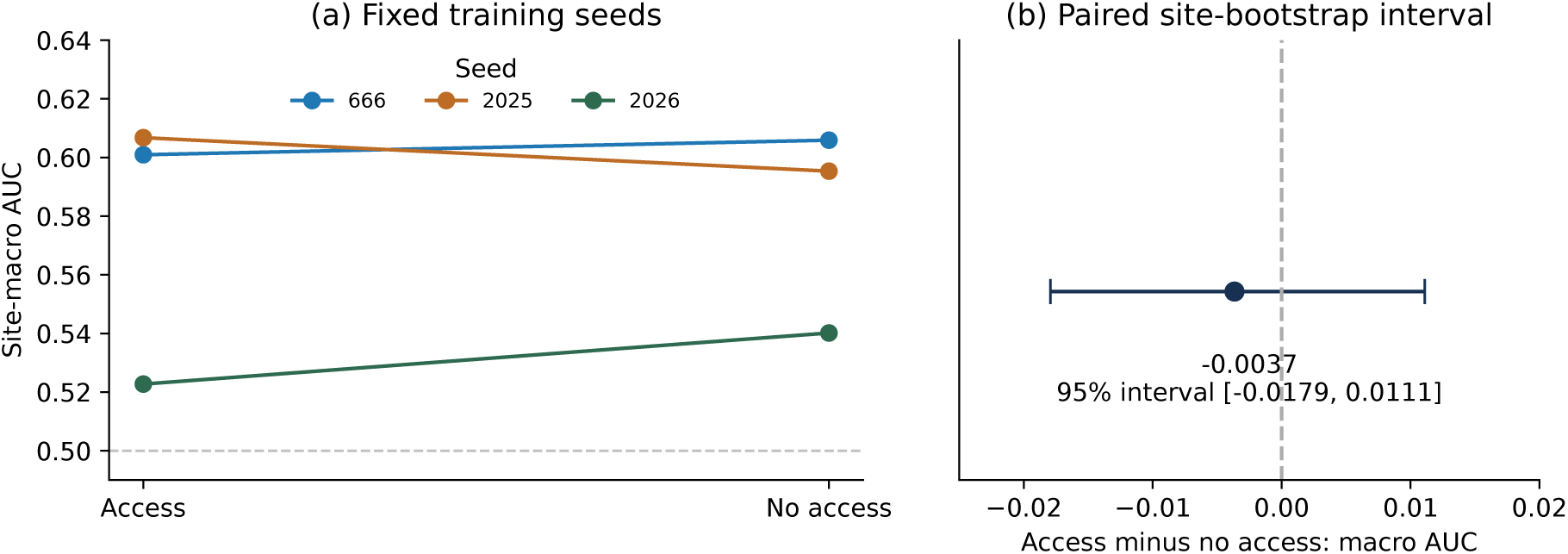
Supplementary scoring-time LOSO access comparison. Each point in the left panel is a seed-specific site-macro AUC over the same 20 sites. The right panel shows the three-seed mean access-minus-no-access site-macro difference and its paired site-bootstrap 95% interval. The training procedure is the same in both conditions; target access edges are inserted only at scoring. Intervals condition on frozen predictions and the three fixed seeds.

The separate site-wise supervision-masking protocol produced pooled AUC 0.480 (0.374–0.584) across 871 subjects. High discrimination was therefore not retained under that intervention, although its difference from cohort 10-fold performance is not a formal paired effect. Exploratory label-proximity and within-site training-label permutation analyses provide further constraints on interpretation (Supplementary Notes S3–S4), with incomplete provenance and public reconstruction limits.

### 4.4. Supplementary unseen-site access comparison

The scoring-time LOSO analysis yielded mean site-macro AUC 0.5768 with population-edge access and 0.5805 without it. The paired access-minus-no-access difference was −0.0037 (95% site-resampling interval [−0.0179, +0.0111]); 29/60 site–seed differences were positive, a descriptive count rather than 60 independent replicates. No clear average scoring-time access effect was observed under this implementation; this is not an equivalence result or a test of the cohort experiment’s same-site supervision mechanism. Per-seed and pooled endpoints are given in Supplementary Note S8. The historical C-PAC/NIAK LOSO values are not used as current quantitative evidence because their prediction-to-run linkage was not re-established in this revision.

### 4.5. Architecture-specific dependence

In the independent head-ablation campaign, the dual-channel reference achieved 0.949 (0.883–0.979); it is not the same fitted run as the cohort reference of 0.941. Single-channel Transformer-Conv, GAT, GCN and no-graph MLP heads yielded 0.554, 0.516, 0.499 and 0.579, respectively (Table 4; Figure 5). The archived head differences are descriptive here: their original paired significance and Holm values have not been re-established from the current evidence package. The single-channel head’s three seed values (0.5544, 0.5983 and 0.5635) also did not reproduce the approximately 0.95 AUC (Supplementary Table S7).

**Figure 5:**
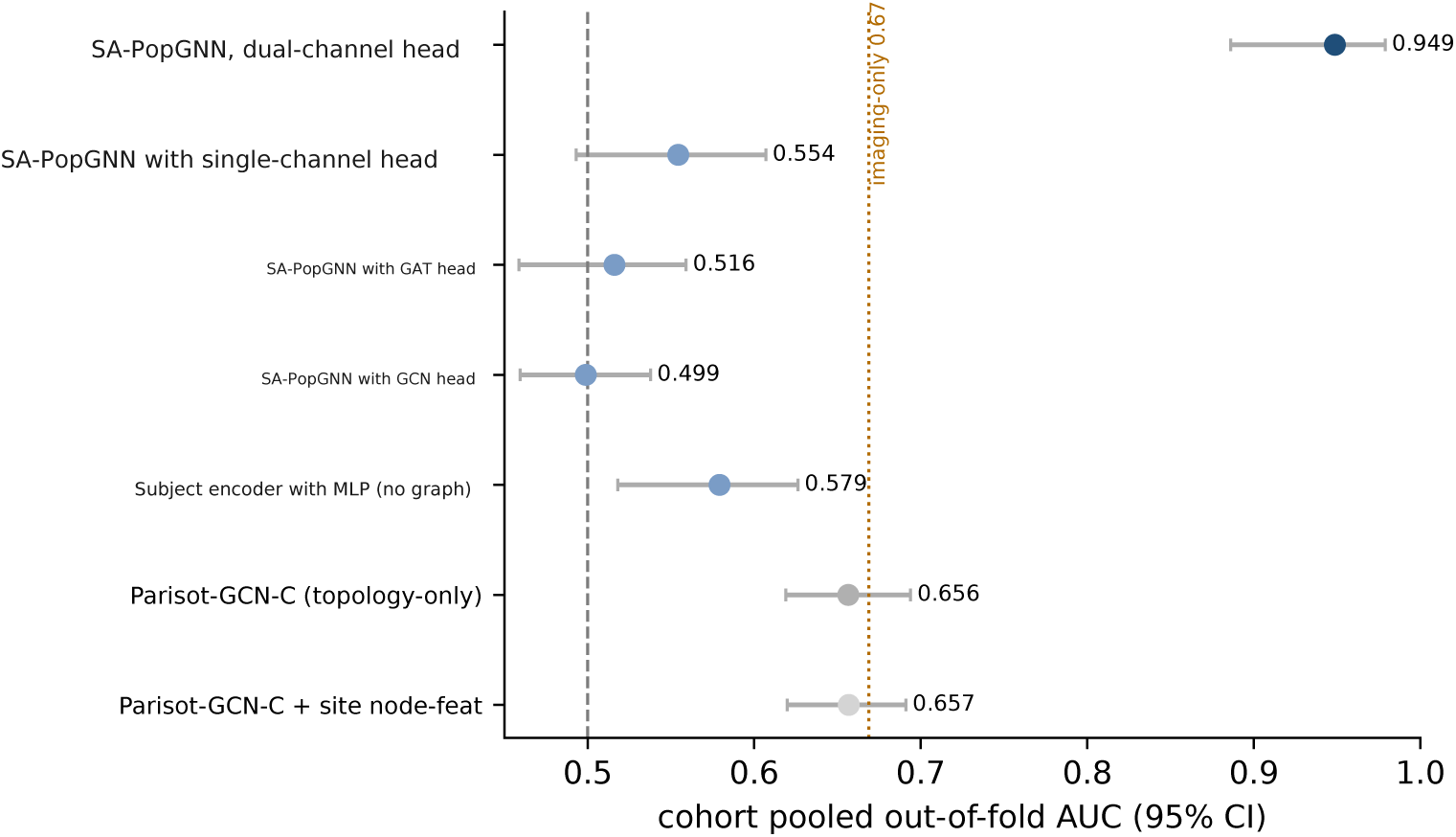
Archived architecture-matched head summaries and topology-only comparator; paired significance is not re-established here. The head ablations retain the SA-PopGNN encoder, sex+site edges, folds and training protocol. In the archived head campaign, the evaluated sex-relation dual-channel head retained approximately 0.95 cohort AUC; all alternative heads yielded lower discrimination. Comparator values come from the separately specified Parisot-GCN-C implementation.

**Table 4:**
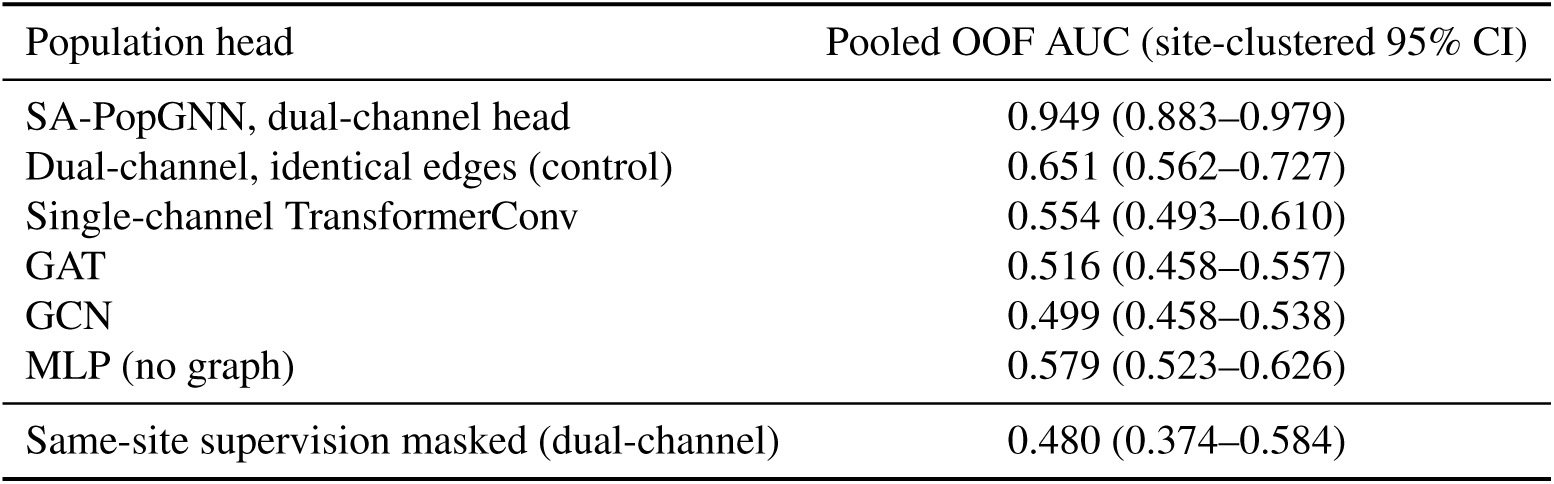
Architecture-matched head ablation under the cohort 10-fold protocol (same SA-PopGNN subject-level connectome encoder, same sex+site edges, same folds and training; only the population head is swapped), followed by the site-wise supervision control (last row); the half-site arms are reported separately in Supplementary Material (Supplementary Note S2) and are not head replacements. Archived pooled OOF AUC and site-clustered intervals; paired significance is not re-established in this revision.

| Population head | Pooled OOF AUC (site-clustered 95% CI) |
| --- | --- |
| SA-PopGNN, dual-channel head | 0.949 (0.883–0.979) |
| Dual-channel, identical edges (control) | 0.651 (0.562–0.727) |
| Single-channel TransformerConv | 0.554 (0.493–0.610) |
| GAT | 0.516 (0.458–0.557) |
| GCN | 0.499 (0.458–0.538) |
| MLP (no graph) | 0.579 (0.523–0.626) |
| Same-site supervision masked (dual-channel) | 0.480 (0.374–0.584) |

**Table 5:** Recomputed paired contrasts from the separately supplied seed-666 control runs (871 subjects, 20 sites). Full AUC is 0.949842 and site-only AUC is 0.955977. Intervals use paired site resampling, 10,000 draws, preserving repeated sites. These runs are not identified as the archived headline model. No historical *p*-value or Holm correction is validated by this table.

| Comparison | $\Delta$ pooled AUC | 95% site interval |
| --- | --- | --- |
| Full vs site-only | −0.0061 | [−0.0234, +0.0096] |
| Full vs sex-only | +0.3800 | [+0.3098, +0.4240] |
| Full vs random | +0.3695 | [+0.3091, +0.4270] |
| Site-only vs random | +0.3756 | [+0.3196, +0.4288] |
| Full vs subject-only | +0.3863 | [+0.3122, +0.4327] |
| Full vs fused-feature shuffle | −0.0111 | [−0.0324, +0.0034] |
| Full vs no train–test edges | +0.3842 | [+0.3434, +0.4372] |
| Site-only vs no train–test edges | +0.3903 | [+0.3525, +0.4373] |
| Site-only vs rewiring | +0.3579 | [+0.3061, +0.4190] |

Supplying identical edges to both branches reduced dual-channel AUC to 0.651 (0.562–0.727). This constrains a two-branch-capacity explanation and implicates relation-specific routing within the tested head family, while leaving graph structure, optimization and head-specific tuning as possible contributors.

The topology-only Parisot-GCN-C comparator showed a different profile: cohort AUC 0.656 (0.619–0.694), site-only 0.650 (0.615–0.688), against a cohort imaging-only reference of 0.660. Its historical LOSO value is retained only in the supplementary provenance record. Adding site as a node feature yielded 0.657/0.658 (Supplementary Table S8). The high cohort pattern was therefore absent in this evaluated comparator. This descriptive cross-architecture comparison does not isolate architecture from optimization and other implementation differences, nor test sensitivity to future cohort-composition shifts.

### 4.6. Paired support-location and implementation sensitivity

The budget-matched paired experiment showed opposite conclusions across edge constructions. With directed edges, the across-site mean ΔAUC was +0.252 (site-resample 95% interval [+0.184, +0.322]), positive at 17/18 sites (exact two-sided sign-test *p* = 1.5 × 10^−4^). With symmetrised edges, it was −0.003 ([−0.020, +0.015]), positive at 8/18 sites. Thus the positive directed effect was not retained after symmetrisation; it is implementation-conditional, not evidence of a general target-site support advantage. The endpoint is the mean of within-site fold AUC differences, averaged equally across 18 eligible sites and three seeds, not a single pooled OOF AUC difference. There are 88 eligible site–fold cells and 836 distinct evaluated subjects; the parent cohort contains 871 subjects from 20 sites. Symmetrisation is a parallel implementation-sensitivity analysis and materially changes the interpretation of the directed result.

Across the three fixed training seeds, directed across-site means were +0.217, +0.256 and +0.283. Site effects and seed variability are shown in Figure 7; signs agreed between constructions at 7/18 sites. The graph, validation and training schedule were matched within pairs (Figure 6), but support location still changed together with subject identity and covariate composition.

**Figure 6:**
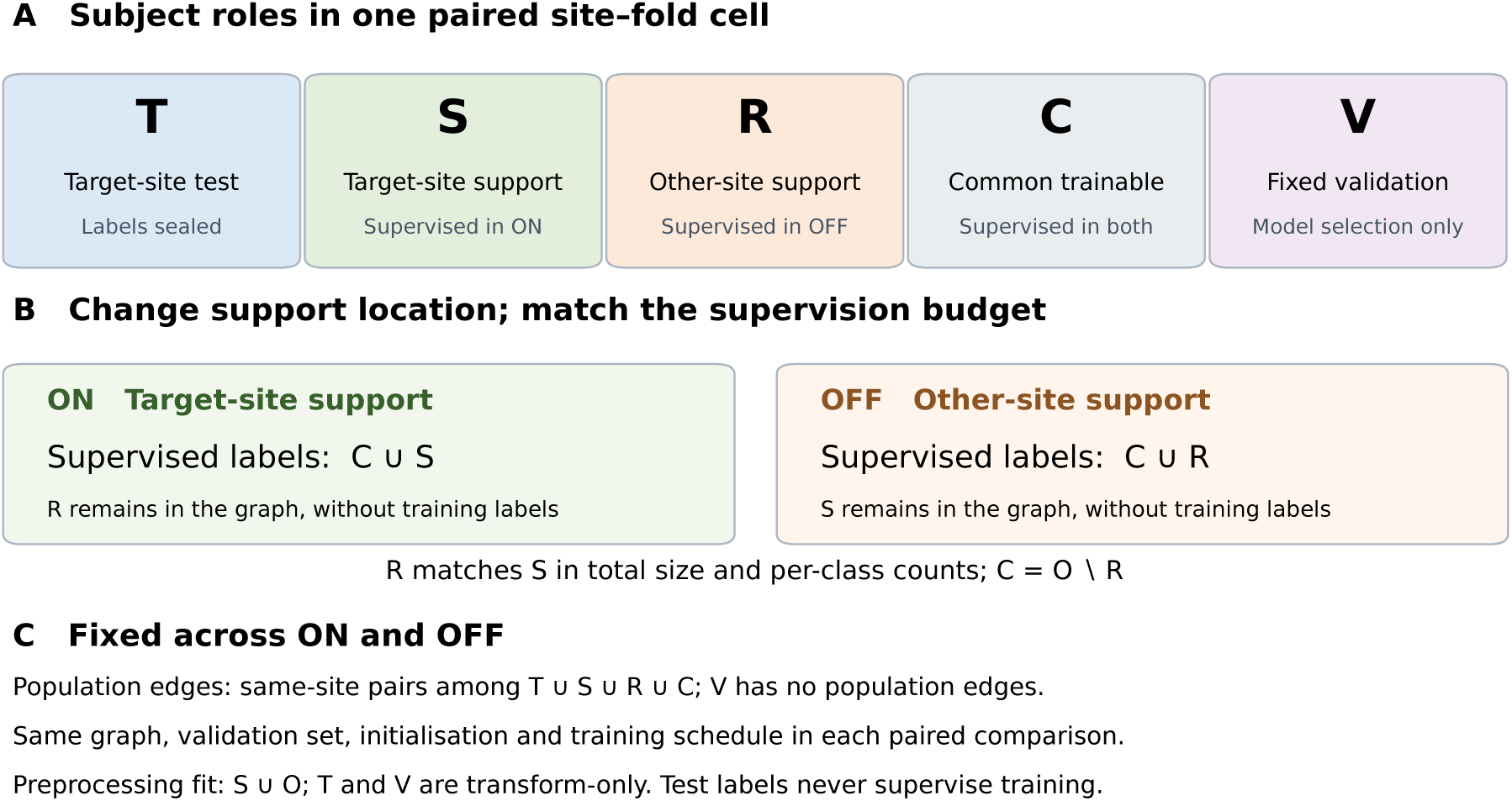
Paired supervision-location protocol. (A) *T* is the target-site test fold, *S* target-site support, *R* other-site support, *C* the common trainable set, and *V* fixed validation. Let *O* denote other-site trainable subjects, excluding *V*; *R* ⊂ *O* matches *S* in total size and per-class counts, and *C* = *O* \ *R*. (B) Training labels are supplied by *C* ∪ *S* in ON and *C* ∪ *R* in OFF; the unsupervised support set remains graph-visible. (C) The graph, validation split, initialisation and training schedule are identical within each pair. Same-site population edges connect nodes in *T* ∪ *S* ∪ *R* ∪ *C*, while *V* has no population edges. Preprocessing is fitted on *S* ∪ *O*; *T* and *V* are transform-only, and test labels remain sealed. The second TransformerConv channel receives no edges but retains its root-weight transform. The primary construction writes each edge pair once; Figure 7 also reports the symmetrised construction. Matching supervision budgets does not isolate support location from subject identity and covariate composition.

**Figure 7:**
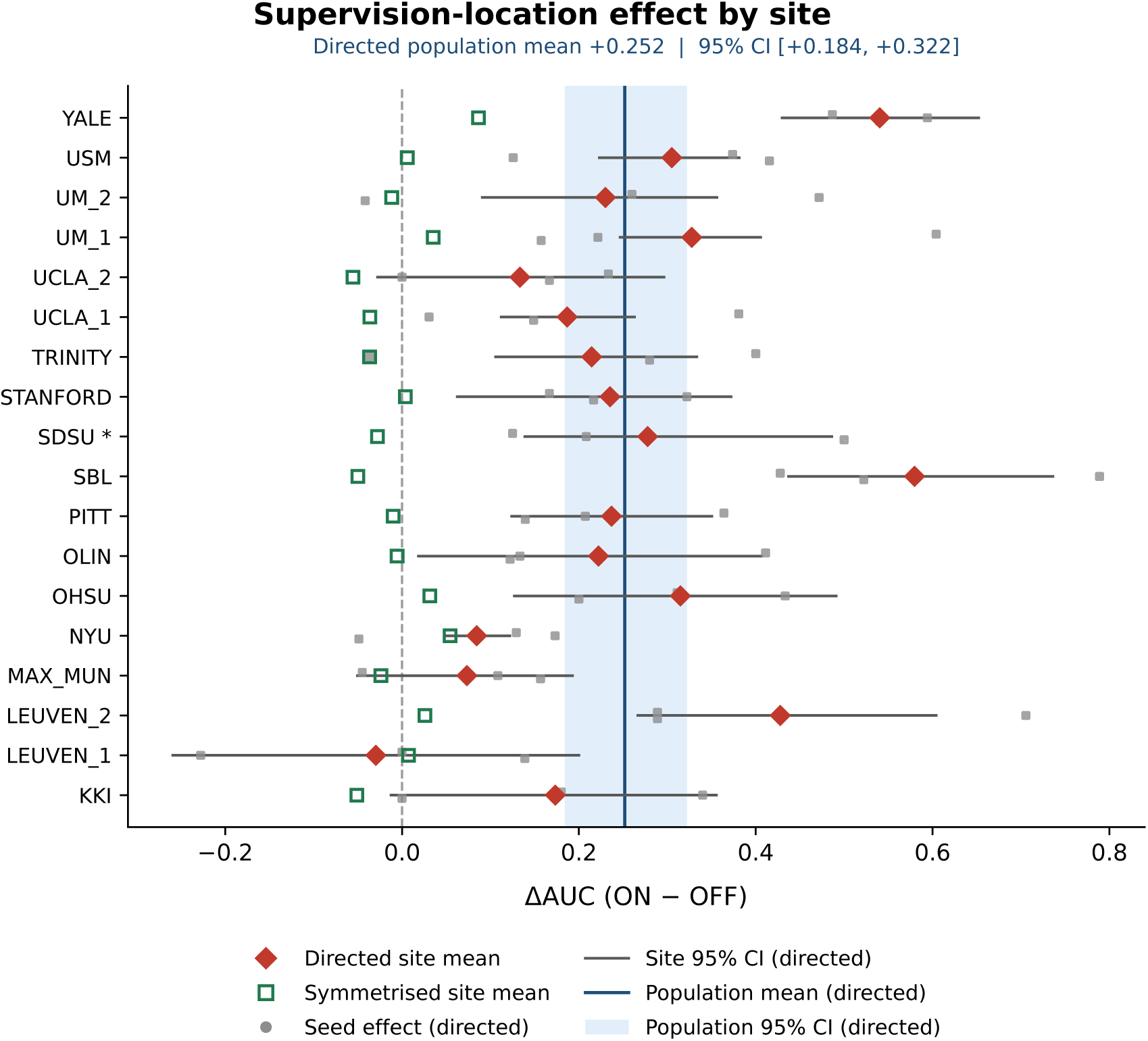
Per-site supervision-location effect under both edge constructions, expressed as ΔAUC (ON minus OFF). Red diamonds and open green squares show site means over three training seeds for the directed and symmetrised constructions, respectively. Grey dots are the three directed seed effects, vertically offset only for readability; horizontal lines are the directed within-site conditional 95% intervals. These intervals condition on fitted models and the fixed seed set; they do not quantify uncertainty from retraining or new training seeds. The blue line and band show the equal-weight directed population mean (+0.252) and site-resample 95% interval ([+0.184, +0.322]), respectively; the band is not a per-site interval. SDSU (asterisk) has three eligible folds; all other displayed sites have five. CALTECH and CMU have no eligible cells and are excluded. The symmetrised population summary is −0.003 ([−0.020, +0.015]), reported as a parallel robustness analysis.

#### Support-draw sensitivity

Across three pre-fixed draws, directed site effects were +0.065/−0.072/− 0.024 at LEUVEN_1, +0.071/ + 0.117/ + 0.007 at PITT and +0.652/ + 0.654/ + 0.572 at YALE. At the first two sites, between-draw variation was comparable to or larger than the effect itself; at YALE the positive effect exceeded that variation. The ON arm also changed across draws in 34/45 comparable cell–seed pairs: the graph was fixed, but its supervised set *C* ∪ *S* changed because *C* = *O* \ *R*. This descriptive sensitivity reinforces the entanglement of location and support composition and is not pooled with the across-site endpoint.

## 5. Discussion

The archived cohort runs show high discrimination under site-aware graph settings, while the newly recomputed controls document related dependencies in a separately identified control campaign. Neither evidence source establishes unseen-site performance or a complete mechanism. The training-only site-prior baseline did not reproduce high cohort AUC, constraining a simple site-prevalence explanation without excluding other site-related pathways.

The paired support-location effect was sensitive to implementation: the positive directed result was absent after symmetrisation. Direction follows subject ordering in that construction, and support resampling changes supervised subject composition. The evidence therefore supports an implementation-dependent observation rather than a stable causal advantage of same-site supervision.

The historical fused-feature shuffle cannot establish dispensability of complete individual imaging, because connectivity topology was retained and feature origins could cross fold roles. Later complete-imaging and added-access campaigns are not used to repair that inference: the former did not match the original graph protocol, and the latter used a new graph construction with split deviations in two seeds.

The supplementary LOSO access result is narrower. Under the same source-site training procedure, insertion of population edges at scoring did not show a clear average site-macro effect. It does not establish equivalent performance, complete information isolation, or the pathway carrying a cohort supervision effect. Descriptive architecture differences likewise concern the evaluated implementations, with optimization and provenance differences unresolved.

### 5.1. Implications for multisite imaging evaluation

Transductive population graphs can be appropriate when deployment genuinely supplies unlabeled target subjects and labeled development subjects under the stated graph contract. A batch of target scans alone, however, does not provide the same-site labeled support available in cohort cross-validation. Authors should distinguish cohort-conditioned performance from unseen-domain performance and specify which target imaging, phenotypes, edges and labels are visible in each regime.

This extends reliability and domain-generalization concerns in medical imaging [1, 2] to models that explicitly use cohort relations. For population graphs incorporating site information [9, 10], useful controls include an imaging-only reference, a site-only graph where applicable, pathway interventions and site-held-out evaluation (Table 6). Prospective single-subject testing is additionally relevant when no target cohort is available. Imaging-only references should be evaluated under compatible site-held-out splits and preprocessing, as in earlier inter-site ABIDE work [22]; they provide context rather than a performance ceiling. Historical imaging-only LOSO figures are not used to rank the new access analysis.

**Table 6:** Suggested reporting items for multisite population-graph evaluation. These items should be selected for the model and intended use; they are methodological recommendations rather than a universal journal requirement.

| Item | Information or question to report |
| --- | --- |
| Information-access contract | Target imaging and phenotype visibility; graph construction; fitted preprocessing; optimization and model-selection labels. |
| Imaging-only reference | Discrimination available without inter-subject graph context, using compatible splits and estimators. |
| Site-only graph, if applicable | Whether site-defined connectivity retains the cohort result; state the remaining head and feature inputs. |
| Train-test pathway control | Which cross-partition edges are removed and whether validation paths remain; distinguish pathway changes from optimization effects. |
| Site-held-out evaluation | Target-site exclusion from fitting and selection; target-node attachment and internal edges; pooled and site-level summaries. |
| Uncertainty and provenance | Resampling unit, seed repetitions, comparison family, saved masks and predictions, and reconstruction limits. |

### 5.2. Limitations

#### Data and generalization

The study uses one dataset, ABIDE-I, and the dependence pattern concerns one site-aware model family. Training variability is covered only by available seed repetitions; not every intervention has three seeds. Small held-out sites produce uncertain site-level estimates. LOSO changes supervision, graph topology and training-domain composition together, and we did not test prospective single-subject prediction or cohort-composition shifts. The demographic baseline covers simple site/sex terms rather than every demographic or acquisition-related pathway.

#### Design and inference

The paired result depends on edge construction and a fixed support draw per cell; supervision location and support composition remain entangled. Its across-site endpoint was implemented post hoc and was not pre-registered. Bootstrap intervals condition on fixed predictions and seeds and do not capture full retraining uncertainty. Non-significant contrasts are not equivalence tests. Some archived analyses have summary evidence only. Historical paired significance and Holm adjustments are retained only for traceability and are not current inferential evidence. Newly recomputed control-run intervals do not recover the identity of historical fits. The LOSO access analysis uses three fixed seeds and a scoring-time intervention; training-time target access and prospective deployment are not tested.

#### Implementation and provenance

Numerical matching verifies the tested model path, not every original training artifact. The NIAK cache builder is unavailable, and zeroed context features cannot be traced to a verified preprocessing stage. The recovered mechanism-control cache lacks a campaign-time hash; its correspondence is supported by subject order, fold/label pairing and other records rather than byte-level proof. The supplementary label-permutation re-run is not fully reconstructible from the public release. Details are consolidated in Supplementary Note S5. Integrated-gradients attribution was insufficiently stable across folds (top-20 ROI overlap 0.23) for anatomical interpretation. No clinical or deployment claim follows from these analyses.

## 6. Conclusion

High transductive cohort AUC in the evaluated site-aware population-GNN is insufficient to establish unseen-site ASD performance. A simple training-only site-prior classifier did not reproduce that high AUC, and the paired support-location result changed substantially with edge construction. The supplementary scoring-time LOSO access comparison showed no clear average effect under its specified implementation. These findings support explicit information-access contracts, compatible estimators, implementation-sensitivity checks and run-level provenance; they do not establish a complete mechanism or a general property of population-GNNs.

## Declarations

### CRediT authorship contribution statement

**Keliang Wan:** Data curation, Investigation, Writing – original draft. **Zongqing Chen:** Conceptualization, Methodology, Investigation, Writing – original draft, Writing – review and editing, Supervision, Project administration, Funding acquisition. **Biantian Yu:** Investigation, Data curation. **Qian Zhang:** Resources, Validation, Supervision (clinical expertise). **Guixian Liu:** Methodology, Formal analysis, Writing – review and editing. **Fanghong Zhang:** Supervision, Funding acquisition, Writing – review and editing. **Ning Zhong:** Supervision, Writing – review and editing. **Hongzhi Kuai:** Writing – review and editing.

### Ethics approval

This study was a secondary analysis of publicly available, de-identified data from the Autism Brain Imaging Data Exchange I (ABIDE-I). Ethics approval for the original data collection was obtained by each contributing site in accordance with its local institutional requirements. No additional ethics approval was required for the present secondary analysis.

### Consent to participate

Written informed consent was obtained by the original contributing sites from adult participants or from the parents or legal guardians of minor participants; assent was obtained where applicable under the local protocols. No participants were newly recruited for this secondary analysis.

### Consent for publication

Not applicable. This study used publicly available, de-identified data and contains no identifiable individual participant information.

### Data and code availability

ABIDE-I data are distributed by the ABIDE consortium; no primary imaging data are redistributed with this article. Code, frozen analysis results and selected recomputation inputs are archived in Zenodo (version 1.2.1, https://doi.org/10.5281/zenodo.23445229), corresponding to GitHub tag `v1.2.1` at commit `84933c3dfc2bdb4b653b3eaacc938cdcba04f540`. The archive includes the existing R3 materials and additional code and aggregate summaries for the analyses reported in this revision. Additional participant-level predictions and reconstructed masks are not publicly released. Consequently, the archive does not support complete individual-level recomputation of all added analyses or end-to-end reconstruction of every historical and supplementary training experiment. Analysis-specific availability and provenance limitations are described in Supplementary Notes S4 and S7–S9.

## Acknowledgments

The numerical calculations in this paper were supported by the Hefei Advanced Computing Center, Hefei, Anhui, China.

## Declaration of generative AI and AI-assisted technologies in the manuscript preparation process

During the preparation of this work, the authors used ChatGPT/Codex (OpenAI) to support language polishing, manuscript organization, and editorial revision, and OpenAI ChatGPT image-generation tools (accessed September–October 2026) to support the initial design and iterative refinement of the explanatory study-design schematic in Figure 1 and its supplementary panel reproductions. The authors are responsible for reviewing, editing, and verifying all text, scientific content, and final vector artwork and for the content of the submitted manuscript. The specific image-generation model/version was not retained in the supplied provenance record; tool access dates and the authors’ reconstruction of the final artwork are disclosed in the relevant captions.

