## Supplementary Material for "Reliable Evaluation of Transductive Population Graph Neural Networks for Multisite Autism fMRI"

### Evidence status in this revision

Tables S1–S10 and Notes S1–S6 reproduce archived material from the baseline manuscript. Cohort, head, calibration and historical LOSO quantities in those sections have not all been re-established from prediction-level inputs in the current revision. In particular, historical LOSO values are not current evidence for unseen-site performance, and historical significance or Holm values are not endorsed as current tests. Notes S7–S9 distinguish newly recomputed numerical evidence, its run identity, and excluded later campaigns. Historical archival material must not be combined with the new control runs as if it came from one fitted run.

Table S1: Separate pipeline-specific site summaries (seed 666). Panel A reports C-PAC in-cohort results and same-site training neighbours; Panel B reports NIAK LOSO results. **Panel B is an unverified historical LOSO record. The panels are not a within-pipeline cohort-versus-LOSO comparison and must not be subtracted.**

**Panel A: C-PAC, in-cohort**

| Site | *n* | ASD | ASD prop. | in-cohort AUC | train neighbours |
|---|---|---|---|---|---|
| NYU | 172 | 74 | 0.43 | 0.996 | 138.4 |
| UM_1 | 86 | 34 | 0.40 | 0.990 | 71.3 |
| USM | 67 | 43 | 0.64 | 0.982 | 52.3 |
| UCLA_1 | 64 | 37 | 0.58 | 0.973 | 54.2 |
| PITT | 50 | 24 | 0.48 | 0.939 | 40.5 |
| MAX_MUN | 46 | 19 | 0.41 | 0.536 | 37.4 |
| TRINITY | 44 | 19 | 0.43 | 1.000 | 34.1 |
| YALE | 41 | 22 | 0.54 | 0.981 | 33.0 |
| UM_2 | 34 | 13 | 0.38 | 0.890 | 26.8 |
| KKI | 33 | 12 | 0.36 | 0.568 | 25.9 |
| LEUVEN_1 | 28 | 14 | 0.50 | 0.531 | 24.5 |
| LEUVEN_2 | 28 | 12 | 0.43 | 0.802 | 21.6 |
| OLIN | 28 | 14 | 0.50 | 0.918 | 21.6 |
| SDSU | 27 | 8 | 0.30 | 0.993 | 18.4 |
| SBL | 26 | 12 | 0.46 | 0.786 | 19.2 |
| OHSU | 25 | 12 | 0.48 | 0.981 | 18.5 |
| STANFORD | 25 | 12 | 0.48 | 0.994 | 19.1 |
| UCLA_2 | 21 | 11 | 0.52 | 0.955 | 15.9 |
| CALTECH | 15 | 5 | 0.33 | 1.000 | 12.3 |
| CMU | 11 | 6 | 0.55 | 0.733 | 8.9 |

**Panel B: NIAK, leave-one-site-out**

| Site | LOSO AUC |
|---|---|
| NYU | 0.541 |
| UM_1 | 0.502 |
| USM | 0.654 |
| UCLA_1 | 0.655 |
| PITT | 0.668 |
| MAX_MUN | 0.593 |
| TRINITY | 0.669 |
| YALE | 0.531 |
| UM_2 | 0.557 |
| KKI | 0.587 |
| LEUVEN_1 | 0.648 |
| LEUVEN_2 | 0.604 |
| OLIN | 0.551 |
| SDSU | 0.566 |
| SBL | 0.280 |
| OHSU | 0.641 |
| STANFORD | 0.462 |
| UCLA_2 | 0.354 |
| CALTECH | 0.560 |
| CMU | 0.567 |

Table S2: Sex-stratified in-cohort performance of the seed-666 model (the small female ASD sample precludes fairness claims).

| Group | $n$ | ASD | AUC (95% CI, subj.) | AUC (95% CI, site-clustered) |
|---|---|---|---|---|
| Male | 727 | 349 | 0.941 (0.924–0.957) | 0.941 (0.869–0.977) |
| Female | 144 | 54 | 0.931 (0.886–0.968) | 0.931 (0.806–0.980) |

Table S3: In-cohort calibration of the seed-666 model. Brier score 0.098; expected calibration error (10 bins) 0.040.

| Metric | Value |
|---|---|
| Brier score | 0.098 |
| Expected calibration error (ECE) | 0.040 |

The Parisot-GCN-C comparator is an independent re-implementation of the published design inside this harness; no upstream code or original dataset artifact was used. Nodes carry reduced subject representations. This comparator uses directed Chebyshev propagation with self-loops and row normalization (one layer, 16 hidden units, order three). That path is distinct from the TransformerConv population head and the paired support-location experiment described in the main text.

Table S4: Estimator comparison for the cohort-level reference values (seed 666). Pooled out-of-fold AUC is the primary estimator for the transductive models; mean-fold AUC is also reported to permit comparison with the mean-fold imaging reference. Mean-fold AUC is recomputed from the frozen outer partition joined by subject ID; pooled and mean-fold AUC weight subjects and folds differently and are reported as distinct estimators.

| Model | pooled OOF AUC | mean-fold AUC | site-clustered 95% CI |
|---|---|---|---|
| Full model (sex+site) | 0.9405 | 0.9466 | 0.867–0.976 |
| Site-only | 0.9480 | 0.9503 | 0.875–0.982 |
| Imaging-only *k*NN (cross-site) | 0.6603 | – | 0.609–0.695 |
| Imaging RF (cohort, mean-fold) | – | 0.6689 | – |

Table S5: Historical leave-one-site-out summary, not revalidated in this revision; seed 666, $n$ = 20 sites, reported as recomputed in the historical record (2026-09-11); the prediction-to-run linkage is unresolved here. The pooled value weights subjects and the macro value weights sites; the site-bootstrap CI resamples sites and includes 0.5, consistent with the absence of clear unseen-site discrimination.

| Pipeline | pooled AUC | site-clustered 95% CI | site-macro AUC | site-median AUC |
|---|---|---|---|---|
| C-PAC | 0.5216 | 0.478–0.558 | 0.5218 | 0.5199 |
| NIAK | 0.5317 | 0.485–0.586 | 0.5595 | 0.5662 |

Table S6: Demographic shortcut baseline (S1), canonical frozen folds, seed 666. Regularised logistic regression with $C$ fixed at 1.0; encodings fitted on the training partition of each fold only; intervals are obtained by site-clustered bootstrap over the 20 sites (2,000 replicates). All levels were close to 0.5, and their site-clustered confidence intervals included 0.5. A simple demographic classifier does not reproduce the cohort AUC reported in the main text. This baseline covers additive site and sex structure and their two-way interaction only; it does not exclude site-typical feature distributions.

| Design | pooled OOF AUC | mean-fold AUC | site-clustered 95% CI |
|---|---|---|---|
| intercept / reference | 0.4957 | 0.5 | 0.4679–0.5187 |
| sex | 0.5076 | 0.5292 | 0.4623–0.5386 |
| site | 0.504 | 0.5153 | 0.4089–0.5505 |
| site + sex | 0.5131 | 0.5163 | 0.4247–0.5553 |
| site + sex + site x sex | 0.5006 | 0.5084 | 0.4077–0.5444 |

Table S7: Seed replication for the key conditions. The full and site-only models retain high cohort AUC across these three seeds, whereas the pathway-removed, supervision-masked and single-channel conditions do not reproduce that level. These rows include different evaluation protocols and are not all paired contrasts.

| Condition | seed 666 | seed 2025 | seed 2026 |
|---|---|---|---|
| Full model (sex+site) | 0.9405 | 0.9367 | 0.9397 |
| Site-only edges | 0.9480 | 0.9508 | 0.9533 |
| No train→test edges | 0.5618 | 0.5644 | 0.6152 |
| Supervision masked | 0.4797 | 0.5205 | 0.4860 |
| Single-channel head | 0.5544 | 0.5983 | 0.5635 |

Table S8: Archived cross-architecture comparison (pooled OOF AUC and historical site-clustered intervals). LOSO rows are historical records, not current evidence.

| Setting | SA-PopGNN (site-aware) | Parisot-GCN-C (topology-only) |
|---|---|---|
| Cohort, full | 0.941 (0.867–0.976) | 0.656 (0.619–0.694) |
| Cohort, site-only | 0.948 (0.875–0.982) | 0.650 (0.615–0.688) |
| Cohort, imaging-only | – | 0.660 (0.624–0.694) |
| LOSO (unseen site) | 0.522 (C-PAC) | 0.650 (0.612–0.685) |
| + site node feature | – | 0.657 / 0.658 |

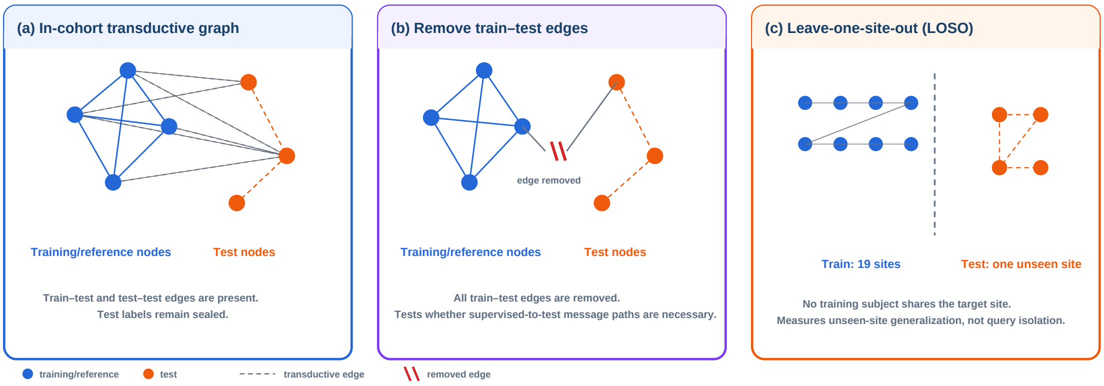


Figure S1: Protocol schematic. (a) In-cohort transduction permits train–test and test–test edges while test labels remain sealed. (b) The train–test edge intervention removes edges linking training to test nodes while preserving within-partition structure. (c) Leave-one-site-out trains on all non-target sites and evaluates one unseen site.

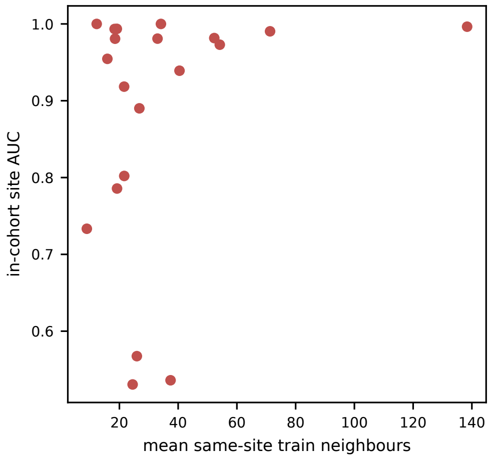


Figure S2: Exploratory association between the mean number of same-site training neighbours and in-cohort site AUC (Spearman $\rho = 0.11$, $p = 0.65$; not significant, descriptive only).

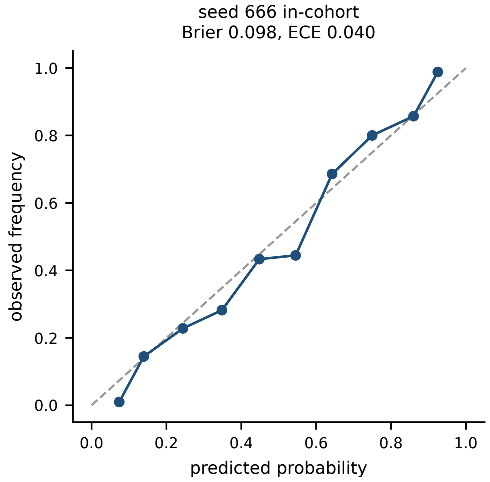


Figure S3: Reliability diagram for the seed-666 in-cohort model (10 bins).

Table S9: Half-site (431-subject) arms, reported descriptively (site-wise protocol; not head replacements and not paired with the cohort 10-fold full model). Pooled out-of-fold AUC, site-clustered 95% CI.

| Half-site arm | Pooled OOF AUC (site-clustered 95% CI) |
|---|---|
| same-site supervision removed, half-site test | 0.474 (0.367–0.594) |
| same-site supervision kept, approximate supervision budget | 0.921 (0.821–0.966) |

**Supplementary Note S1: excluded historical cross-protocol contrast**

The historical paired site-clustered bootstrap analysis was originally computed over 13 contrasts. One of them compared the cohort 10-fold full model with the same-site supervision-masked arm. Both arms use the same 871 subjects, but they were produced under different evaluation protocols (the cohort 10-fold protocol versus a site-wise supervision-masked protocol evaluated over the same 871 subjects), and the identity of the outer partition of the masked arm is not recoverable from the released artefacts. That contrast is therefore not presented as a formal paired comparison. Its archived values are $\Delta$AUC = +0.461 with site-clustered 95% CI $[0.345, 0.557]$, unadjusted paired-bootstrap $p < 0.001$ and Holm-adjusted 0.009 over the original 13-contrast family; the archived randomization analysis reported an unadjusted paired site-level permutation $p = 10^{-4}$. The original table comprised 12 within-protocol contrasts; its archived quantities are not endorsed in the revised main table. Original Holm values refer to the 13-contrast family and are not recalculated. Unverified paired intervals and $p$-values for the five descriptive contrasts are recorded in Supplementary Note S6. The paired-bootstrap $p$-values are floored at $1/1500$ and the randomization $p$-values at $1/10000$, as implemented in the archived scripts. These values are retained solely to document the omitted contrast, not as inferential evidence. The historical statistical inputs were not recorded with file hashes, so their byte-level identity cannot be verified from the available archive. This 871-subject comparison is distinct from the separate half-site analysis reported in Supplementary Note S2.

**Supplementary Note S2: half-site (431-subject) descriptive analysis**

This note reports, in full and descriptively, the half-site analysis referenced in the main text. It is not a paired comparison with the cohort 10-fold full model and is not used to attribute the observed effect to same-site supervision alone.

A separate half-site analysis reported AUCs of 0.474 [0.367–0.594] without same-site supervision and 0.921 [0.821–0.966] with same-site supervision retained, with mean training-set sizes of 745.15 and 754.70, respectively. Both arms were designed to evaluate the same half-site subset; the archived outputs and report are consistent with a shared subset of 431 subjects. The exact historical split and individual-level evaluation mask were not retained. These results are therefore reported descriptively and do not isolate the effect of same-site supervision from all differences in supervision budget or training composition.

### B Transductive information-access contract

#### Cohort-visible graph; test labels sealed

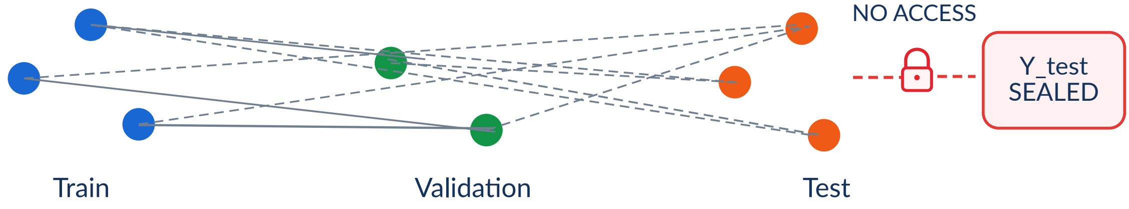


| Information | Train | Validation | Test |
|---|---|---|---|
| Imaging features | YES | YES | YES |
| Site / sex metadata | YES | YES | YES |
| ASD / TD labels | YES | selection only | SEALED |
| Scaler / PCA | fit | transform | transform |
| Train-test edges | YES | YES | YES |
| Test-test edges (in-cohort) | transductive | transductive | transductive |

**Key distinction:** transductive context visibility is permitted; test-label access is not.
Test-test edges exist only in the in-cohort protocol.

Figure S4: Panel B of the study-design figure, reproduced at full size. (B) Transductive information-access contract. Test labels are sealed, while imaging features, site/sex metadata, and cohort graph structure (including test-test edges) are visible in the in-cohort protocol. **AI-assisted illustration:** OpenAI ChatGPT image-generation tools (accessed September–October 2026; specific model/version not retained) were used to support the initial graphical drafting of this explanatory schematic. The authors subsequently reconstructed, edited, and verified the final vector artwork and all scientific content.

#### Supplementary Note S3: exploratory label-proximity analysis (post hoc)

This note reports an *exploratory* analysis that was computed after the frozen campaign, from released artefacts only; it is not part of the frozen result set, it did not influence any number reported in the main text, and it should not be read as a mechanism demonstration in either direction.

*Question and construction.* For each out-of-fold test subject we asked how strongly the predicted ASD probability tracks a simple label-structure quantity available to the model: the fraction of that subject's labelled same-site training neighbours whose label is ASD. The fraction was computed over the fold's outer-training complement only, so no test label can enter it. This complement includes the inner-validation subjects, whose labels are excluded from the optimization loss under

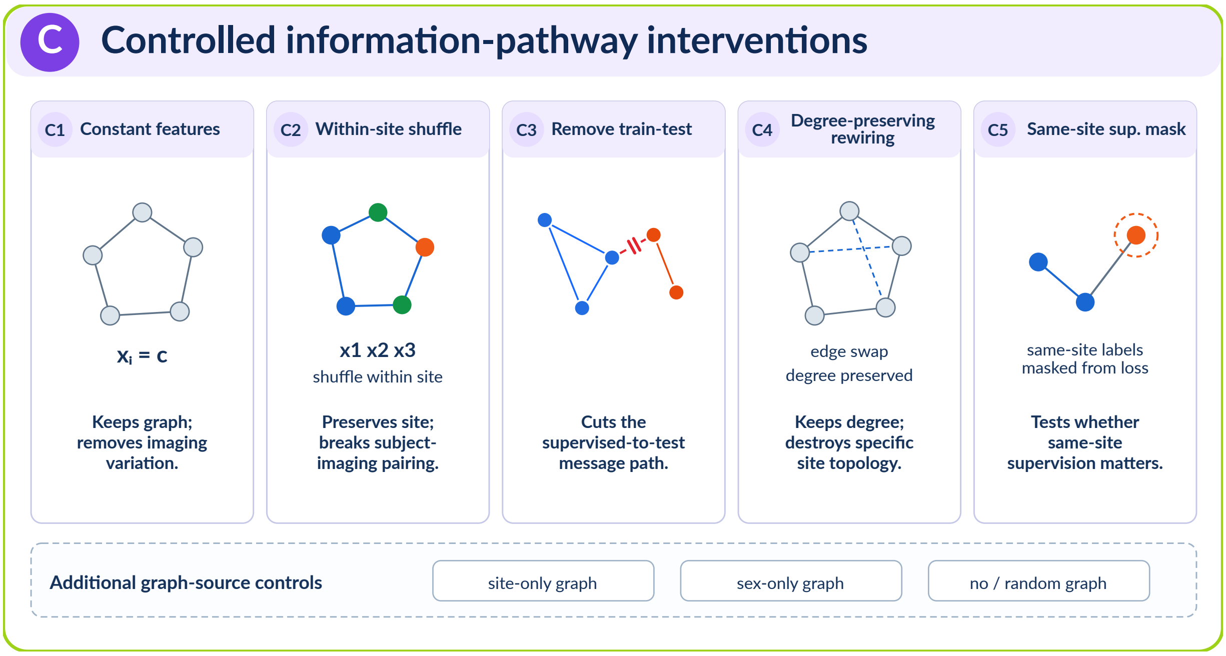


Figure S5: Panel C of the study-design figure, reproduced at full size. (C) Controlled information-pathway interventions and additional graph-source controls used to probe the sources of cohort-level discrimination. **AI-assisted illustration:** OpenAI ChatGPT image-generation tools (accessed September–October 2026; specific model/version not retained) were used to support the initial graphical drafting of this explanatory schematic. The authors subsequently reconstructed, edited, and verified the final vector artwork and all scientific content.

the stated contract; it is therefore an exploratory outer-training label-proximity proxy, not an exact reconstruction of the labels used for supervised optimization. The association uses the released predicted probabilities and this proxy; no model is refitted. Inputs were the released out-of-fold prediction artifact and the hash-pinned phenotype table; the per-subject fold assignment carried by the released prediction artifact was used, and the pooled out-of-fold AUC recomputed from that artifact is 0.9405, matching the value reported in the main text.

*Result.* Across 871 subjects the Spearman correlation between the same-site labelled training-neighbour ASD fraction and the predicted ASD probability was $\rho = 0.1131$; a site-clustered bootstrap (2,000 draws over the 20 sites) gave a 95% interval of [−0.0420, 0.2144], i.e. a weak positive association whose interval includes zero.

*Interpretation (restricted).* A weak association of this kind does not indicate that simple same-site labelled-neighbour prevalence accounts for the observed cohort-level scores. It also cannot exclude that label structure is used in a non-linear or graph-smoothing form, so the interpretation remains restricted to the evaluated interventions and their stated design limitations.

#### Supplementary Note S4: within-site training-label permutation null (E-M1a)

*What was permuted.* Within each training fold the labels of the *training* subset were permuted among subjects of the same site; the validation subset's labels and the test labels were not permuted

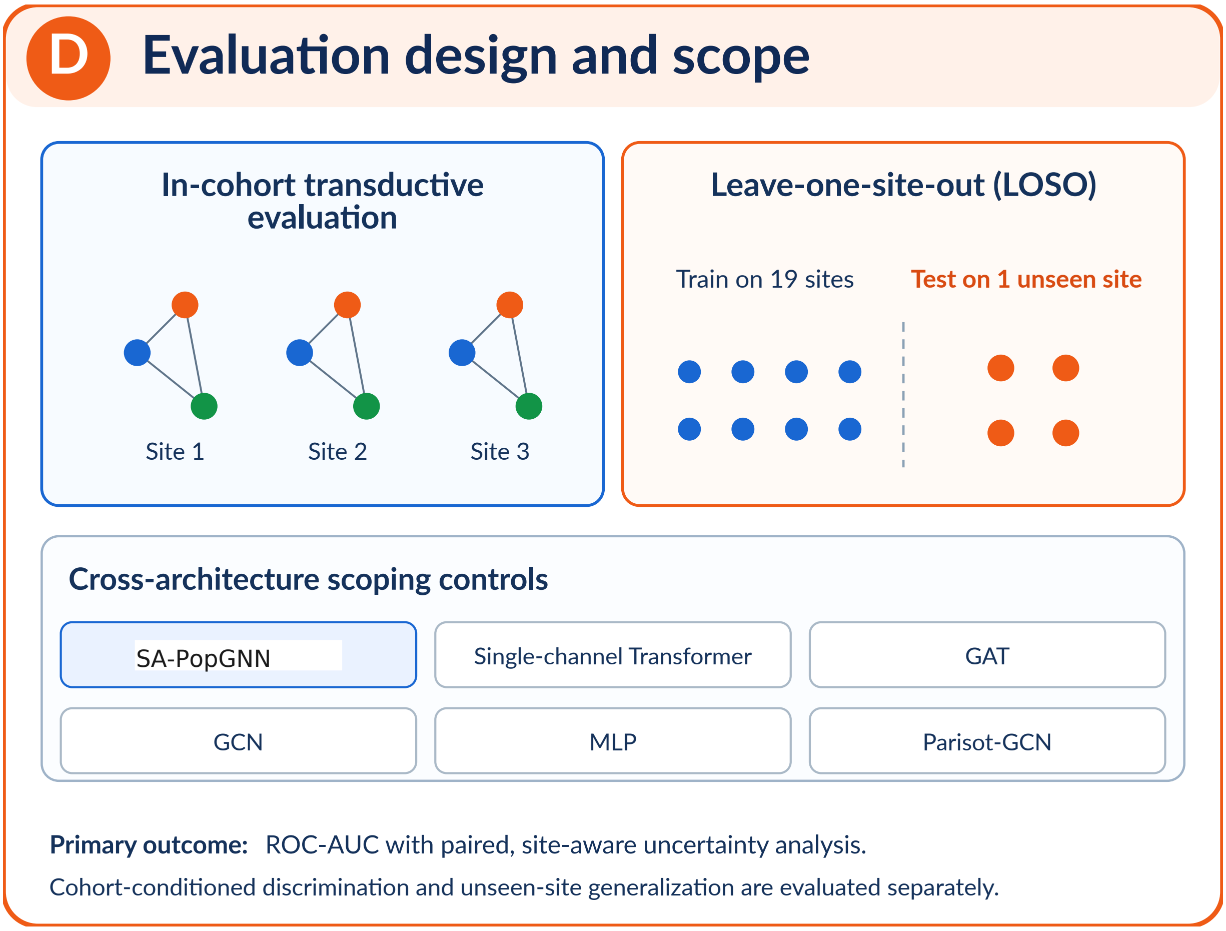


Figure S6: Panel D of the study-design figure, reproduced at full size. (D) Evaluation design and scope, including in-cohort transductive evaluation, leave-one-site-out (LOSO) generalization to an unseen site, and cross-architecture scoping controls. **AI-assisted illustration:** OpenAI ChatGPT image-generation tools (accessed September–October 2026; specific model/version not retained) were used to support the initial graphical drafting of this explanatory schematic. The authors subsequently reconstructed, edited, and verified the final vector artwork and all scientific content.

(asserted at run time: the permutation never touched a test index). The site-only graph, the within-site-feature-shuffle condition, the loss, the optimiser and the fold definition were kept identical to the intervention reported in Table 2, so the intervention is at the level of the training labels that enter the loss; the validation-subset labels were left true and could still influence model selection, and the test labels were never read.

*Protocol identity.* The driver was derived from the campaign's mechanism-control script by a documented, minimal patch (source script sha256 recorded in the run provenance), and was first run *without* permutation: it reproduced the observed within-site-feature-shuffle condition at pooled out-of-fold AUC 0.9627 (reference 0.9604), which supports comparability of the null with the observed value; it does not establish identity of artifacts.

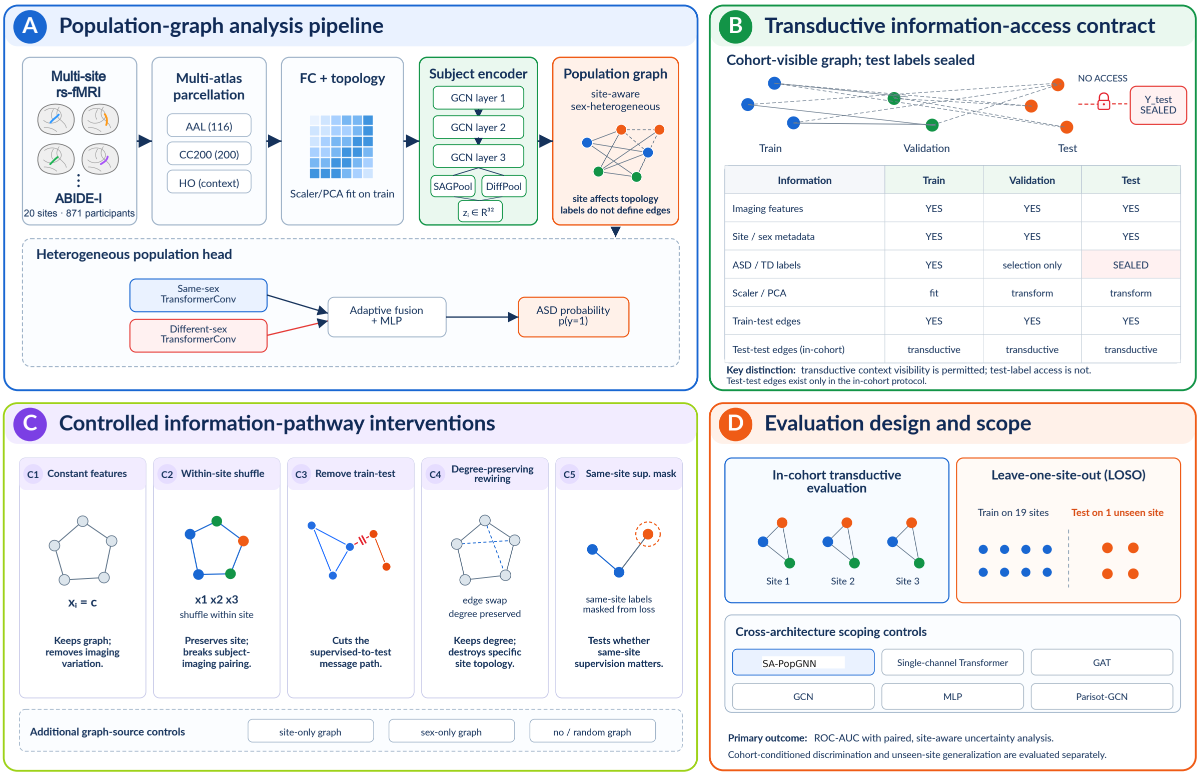


Figure S7: Overview of the analysis pipeline, information-access contract, controlled mechanism interventions, and evaluation design. (A) Multisite resting-state fMRI inputs from ABIDE-I are represented schematically across acquisition sites, followed by multi-atlas parcellation, functional-connectome construction, fold-local scaler/PCA fitting, subject encoding, and population-graph inference. The four brain icons are representative site exemplars rather than four literal sites; the ellipsis indicates additional acquisition sites in the 20-site cohort. Subject embeddings are combined in the site-aware SA-PopGNN population graph, and a dual-channel head with separate same-sex and different-sex branches produces ASD probability. (B) Transductive information-access contract. Test labels are sealed, while imaging features, site/sex metadata, and cohort graph structure (including test-test edges) are visible in the in-cohort protocol. (C) Controlled information-pathway interventions and additional graph-source controls used to probe the sources of cohort-level discrimination. (D) Evaluation design and scope, including in-cohort transductive evaluation, leave-one-site-out (LOSO) generalization to an unseen site, and cross-architecture scoping controls. **AI-assisted illustration:** OpenAI ChatGPT image-generation tools (accessed September–October 2026; specific model/version not retained) were used to support the initial graphical drafting of this explanatory schematic. The authors subsequently reconstructed, edited, and verified the final vector artwork and all scientific content.

*Null distribution.* Across 20 independent permutation seeds the pooled out-of-fold AUC had median 0.4997, mean 0.5014, 2.5–97.5 percentile interval [0.4844, 0.5217] and range [0.4843, 0.5257]. Per-seed values: 1: 0.5173, 2: 0.4930, 3: 0.5170, 4: 0.5021, 5: 0.5000, 6: 0.5257, 7: 0.5156, 8: 0.5145, 9: 0.5092, 10: 0.4944, 11: 0.4843, 12: 0.4880, 13: 0.4910, 14: 0.4988, 15: 0.4845, 16: 0.4993, 17: 0.4989, 18: 0.4890, 19: 0.5015, 20: 0.5035. The observed condition (0.9604) lies above every permuted run. With the finite-sample correction used elsewhere in this work, $(0+1)/(20+1) = 1/21$, i.e. a one-sided tail estimate of approximately 0.0476 (written 0.048); this value is Monte Carlo resolution-limited at 20 permutation seeds and is not a high-precision or

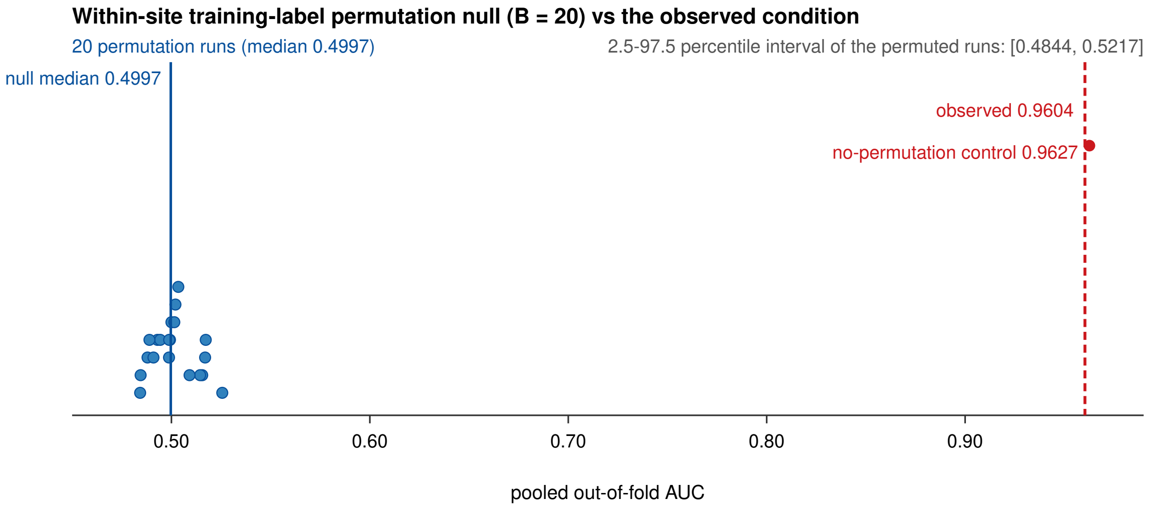


Figure S8: Within-site training-label permutation null (Supplementary Note S4). Each blue point is the pooled out-of-fold AUC of one of the 20 independent permutation runs, in which the labels of the *training* subset were permuted among subjects of the same site; the solid blue line marks the median of the permuted runs. The dashed red line marks the observed release condition (0.9604) and the red point the no-permutation control run of the same driver (0.9627). Validation-subset labels were kept true (they could still influence model selection) and test labels were never permuted. The shaded-free interval quoted in the text is the empirical 2.5–97.5 percentile range of the 20 permuted runs; it is a range of permutation results, not a confidence interval for a population AUC, and the one-sided tail estimate is limited by the Monte Carlo resolution of 20 seeds.

stable significance statement. Because no-permutation training randomness and the provenance limits also affect comparability, this tail estimate is treated as exploratory rather than as a calibrated confirmatory randomization test. The reported 2.5–97.5 percentile interval is the empirical range of the permuted results, not a confidence interval for a population value of AUC. The distribution is shown in Fig. S8.

*Reading of the result (restricted).* The evaluated condition therefore does not survive destruction of the pairing between training subjects and their training labels within site, while the historical fused-feature reassignment retained high AUC with each subject's original connectivity topology preserved. Complete subject-to-imaging correspondence was not tested by that historical control. This is consistent with dependence on training-label pairing in a site-conditioned graph under the evaluated transductive protocol; it does not establish that the model performs label propagation only, and it does not license any statement about unseen sites. The evidence behind this note is not publicly archived: the retained analysis record (an internal record, not a public attachment) contains the driver script, the per-seed tables, the per-run out-of-fold arrays and the recorded run provenance, but not the feature cache itself or a full dependency lock, so this experiment cannot be rebuilt from the released repository alone.

### Supplementary Note S5: implementation and provenance details

*Cache correspondence.* In the available NIAK cache, 10 of 25 context-feature columns are identically zero and 15 are populated. The cache-building script is absent from the audited tree, so the stage at which zeroing occurred is unverified. The cache used by the frozen mechanism controls was recovered from a historical worktree after having been rebuilt from canonical matrices. No campaign-time hash pins its identity. Correspondence is supported by the campaign script's recorded path, the 871-subject order and the fold/label pairing, not by byte-level identity.

The independent Supplementary Note S4 re-run on that rebuilt cache reproduced the no-permutation reference closely (0.9627 versus 0.9604). This supports comparability, not artifact identity. Cross-pipeline direction was also consistent for train–test edge removal (0.562 in C-PAC versus 0.593 in NIAK) without resolving cache provenance. The older LOSO summaries (0.522 versus 0.532) remain historical, unverified records here. The public core release does not by itself make every supplementary analysis reconstructible.

*Paired graph orientation.* The paired experiment records each population edge once with $i < j$, yielding directed propagation, a TransformerConv root-weight transform, no graph self-loops and no row normalization. The second channel has no edges but retains its root transform. A pre-fixed orientation diagnostic changed five of six cell–arm AUCs, by up to 0.250. These six units concern this paired implementation only; they are not diagnostics of the separately constructed main-model or head-ablation graphs. The complete symmetrised parallel analysis, rather than this small diagnostic, supplies the main implementation-sensitivity result.

### Supplementary Note S6: historical paired inference records

The five rows below preserve previously reported paired intervals and significance values whose original statistical inputs could not be relocated. They have not been re-estimated or independently verified in this revision and must not be used as formal inferential evidence. Their point estimates belong to the archived campaign and are not the new control-run table in the main text. The original 13-contrast family is unchanged; no reduced-family Holm correction is calculated.

Table S10: Unverified historical paired quantities, retained for traceability only. The rounded −0.000 endpoint is not an exact zero. These are archived claims, not newly computed or endorsed tests.

| Historical comparison | ΔAUC | Archived 95% interval | Archived $p$ | Archived Holm |
|---|---|---|---|---|
| Full vs subject-only | +0.353 | [0.286, 0.400] | < 0.001 | 0.009 |
| Full vs within-site shuffle | −0.020 | [−0.048, −0.000] | 0.049 | 0.099 |
| Full vs no train–test edges | +0.379 | [0.325, 0.426] | < 0.001 | 0.009 |
| Site-only vs no train–test edges | +0.386 | [0.337, 0.428] | < 0.001 | 0.009 |
| Site-only vs rewiring | +0.351 | [0.300, 0.402] | < 0.001 | 0.009 |

An archived, unadjusted site-level randomization analysis with 10,000 permutations reported $p = 0.48$ for full versus site-only, and $p = 10^{-4}$ for full versus no train–test edges, site-only versus rewiring and dual- versus single-channel heads. These historical sensitivity values are documented without new verification and do not restore formal evidence for contrasts with missing inputs. The separate cross-protocol full-versus-supervision-masked contrast is retained only in Supplementary Note S1.

### Supplementary Note S7: reconstructed empirical site-prior baselines

The main baseline uses all 871 subjects (403 ASD, 468 TD) and 20 sites. Outer folds use StratifiedKFold(10, shuffle=True, random_state=666). For each outer-training partition, validation size is $\max(1, \lfloor 0.1n \rfloor)$ with StratifiedShuffleSplit and random_state=666. Saved inner masks were unavailable; these masks are reconstructed, and the outer folds reproduce the supplied OOF identifiers. For each test subject the score is the ASD fraction among same-site optimization-training subjects. No validation or test label enters this score. There were zero fallbacks. Pooled AUC is 0.5056228924, with a 10,000-draw site-bootstrap percentile interval [0.4176388002, 0.5464389247] (seed 20261015; sites sampled with multiplicity). This interval conditions on fixed scores, not retrained models. Site class counts are in Table S1.

The associated main-model OOF archive has SHA256 beginning `28480a12de230996` (full hash in the retained input manifest, an internal analysis record); its softmax pooled AUC is 0.9396089160 and mean-fold AUC is 0.9424602897. These values are not treated as proof that this archive generated the baseline manuscript's 0.9405. The control-run Full and site-only archives used by the revised main contrast table yield 0.9498419970 and 0.9559765434, respectively. Their input hashes are supplied separately. No historical significance value is transferred to these runs.

An optional empirical-prior baseline for the paired support-location protocol is a separate analysis: ON uses $C \cup S$ labels, OFF uses $C \cup R$. The 836 evaluated subjects contribute one prediction per arm across 88 feasible cells (18 sites); 1,672 rows do not represent independent subjects. ON pooled AUC is 0.5808985857. OFF has no target-site supervised labels and uses the cell-wide optimization-training prior for all 836 predictions, yielding pooled AUC 0.4402782569. Both arms are constant within each cell, so within-cell AUC is 0.5; constants can differ between folds. OFF has site-macro AUC 0.5 and ON 0.4762825928. The pooled OFF value mixes cell-specific constants and is not evidence of a reverse predictive mechanism. No pooled-across-arms AUC is used.

### Supplementary Note S8: scoring-time LOSO population-edge access

This analysis contains 20 held-out sites, three training seeds and two scoring conditions. For every held-out site, source optimization-training and validation sets are shared across conditions; the target site is excluded from both. The source split seed is 666. Scaler/PCA fitting uses optimization-training source subjects only. Training uses the source graph in both conditions, up to 300 epochs, patience 150, with validation accuracy plus AUC for model selection. The target embeddings are concatenated with source embeddings only for scoring. ON inserts cross-site same-sex access edges; OFF omits that insertion. Neither condition has target–target population edges. The second relation channel is passed an empty edge tensor in the supplied driver; architecture-provided root transforms are not equivalent to zero node computation.

The arm switch occurs after source training. Although the driver repeats training per condition with matched seeds, this is not evidence for an effect of training-time target access, and checkpoint equality across all units is not independently established by the supplied predictions. Invariant summaries and deterministic checks are execution records; the local CPU verification independently reproduces the numerical endpoints from the unified prediction table. The external subject-encoder module and LOSO helper are not bundled, limiting full training reconstruction.

Table S11: LOSO access results, independently recomputed from test predictions. Macro AUC gives each of the same 20 sites equal weight; pooled AUC scores all 871 subjects. The mean row averages seed-specific endpoints, not stacked predictions.

| Seed | ON macro | OFF macro | ON pooled | OFF pooled |
|---|---|---|---|---|
| 666 | 0.600938 | 0.605904 | 0.551706 | 0.574961 |
| 2025 | 0.606761 | 0.595341 | 0.584685 | 0.567252 |
| 2026 | 0.522739 | 0.540161 | 0.510005 | 0.553482 |
| Mean | 0.576813 | 0.580468 | 0.548799 | 0.565232 |

For each site and seed, ON minus OFF site AUC is calculated; the endpoint averages sites and then the three fixed seeds equally. Its value is $-0.0036556804$, with percentile interval $[-0.0179319, +0.0111090]$ from 5,000 site draws (seed 20261016). Each draw is shared by both conditions and all seeds. Positive differences occur in 29/60 site–seed entries; these are not 60 independent replicates. A bootstrap positive-draw share is not a $p$-value. No equivalence test or multiplicity-adjusted claim is made. The interval excludes uncertainty from new training seeds and refitting. The earlier 0.5710 summary stacked seed predictions within sites before scoring; it is a different estimand and is not the primary endpoint here.

#### Supplementary Note S9: excluded campaigns and evidence adoption

A later complete-imaging-swap campaign is not used to support dispensability of individual imaging. Its population graph did not correspond to the original cohort graph and an observed response to perturbing other subjects was not localized to a specific computational path. A change in logits alone neither proves a change in class probability nor establishes an abnormal encoder dependency. The campaign is retained as an internal diagnostic record, without further training in this revision.

The added-access endpoint-swap campaign is not used to explain the historical train–test removal result: it introduces edges under a new protocol, and two seeds used the wrong planned folds. The seed-666 exploratory comparison is not part of the manuscript's evidence chain; the deviating units are not pooled into a formal result. The scientific endpoint of this revision is evidence selection, not successful acceptance of every attempted experiment.

The archived directed paired support-location endpoint remains a separate case-study analysis over 18 eligible sites and 88 cells, with 836 distinct tested subjects. Its across-site mean is +0.251853 [0.184089, 0.322206], and symmetrisation yields approximately -0.0032 [-0.0198, 0.0150]. These are means of site-level fold differences, not pooled OOF effects; the change after symmetrisation is substantial implementation sensitivity. The original 13-item Holm family must not be extended to the supplementary analyses.
